# Neutron crystallographic refinement with *REFMAC*5 of the *CCP*4 suite

**DOI:** 10.1101/2023.08.13.552925

**Authors:** Lucrezia Catapano, Fei Long, Keitaro Yamashita, Robert A. Nicholls, Roberto A. Steiner, Garib N. Murshudov

## Abstract

Hydrogen (H) atoms are abundant in macromolecules and often play critical roles in enzyme catalysis, ligand recognition processes, and protein-protein interactions. However, their direct visualisation by diffraction techniques is challenging. Macromolecular X-ray crystallography affords the localisation of the most ordered H atoms at (sub-)atomic resolution (around 1.2 Å or higher), that is not often attainable. Differently, neutron diffraction methods enable the visualisation of most H atoms, typically in the form of deuterium (D) atoms at much more common resolution values (better than 2.5 Å). Thus, neutron crystallography, although technically demanding, is often the method of choice when direct information on protonation states is sought. *REFMAC*5 of the Collaborative Computational Project No. 4 (*CCP*4) is a program for the refinement of macromolecular models against X-ray crystallographic and cryo-EM data. This contribution describes its extension to include the refinement of structural models obtained from neutron crystallographic data. Stereochemical restraints with accurate bond distances between H atoms and their parent atom nuclei are now part of the *CCP*4 Monomer Library, the source of prior chemical information used in refinement. One new feature for neutron data analysis in *REFMAC*5 is the refinement of the protium/deuterium (^1^H/D) fraction. This parameter describes the relative ^1^H/D contribution to neutron scattering for H atoms. The newly developed *REFMAC5* algorithms were tested by performing the (re-)refinement of several entries available in the PDB and of one novel structure (FutA) by using either (*i*) neutron data-only or (*ii*) neutron data supplemented by external restraints to a reference X-ray crystallographic structure. Re-refinement with *REFMAC5* afforded models characterised by *R*-factor values that are consistent with, and in some cases better than, the originally deposited values. The use of external reference structure restraints during refinement has been observed to be a valuable strategy especially for structures at medium-low resolution.

**Synopsis:** The macromolecular refinement package *REFMAC*5 of the *CCP*4 suite has been extended with the incorporation of algorithms for neutron crystallography.

## 1. Introduction

Knowledge of protonation states and hydrogen (H) atom positions in macromolecules can be critical in helping to formulate functional hypotheses and, generally, in providing a more complete characterisation of the biological processes under investigation. H atoms are responsible for the reversible protonation of active site residues involved in enzymatic reactions (Ahmed *et al*., 2007; Fisher *et al*., 2012; Wan *et al*., 2015). They are also necessary for the formation of H-bonds that stabilise macromolecular structures, contributing to the establishment of biological interfaces (Engler *et al*., 2003; Niimura *et al*., 2004; Oksanen *et al*., 2017). Additionally, as H atoms are often involved in determining specificities in protein-ligand recognition processes, their identification and localisation may help the development and design of new therapeutics (Combs *et al*., 2020; Kovalevsky *et al*., 2020; Kneller *et al*., 2022).

The position of many H atoms in macromolecules can be estimated using the coordinates of their parent atoms (those to which they are covalently bound) and known geometric properties (Sheldrick & Schneider, 1997). This is the case, for example, for amide H atoms in the protein backbone, for those bound to C^α^ atoms, for those attached to aromatic carbon atoms, etc. However, many H atoms of biochemical interest, e.g. those on the side chain of histidine, protonated aspartate and glutamate, or those associated with multiple favourable positions (hydroxyl groups of serine, threonine, and tyrosine amino acids) cannot be located on the basis of simple geometric considerations, but need to be determined experimentally (Fisher *et al*., 2009; Gardberg *et al*., 2010).

Although H atoms represent a large fraction of the total atomic content of macromolecules (∼50% and 35% of protein and nucleic acid atoms, respectively) their experimental visualisation is not straightforward. In X-ray macromolecular crystallography they contribute little to the total scattering, thus even at (sub-)atomic resolution (<1.2 Å) only a fraction of all H atoms is typically observed in electron density maps (Howard *et al*., 2004; Petrova & Podjarny, 2004). For instance, in the case of 0.85 Å resolution room temperature X-ray structure of crambin, less than 50% of all H atoms could be identified (Chen *et al*., 2012). These tend to be the most ordered ones, seldom interesting from a functional viewpoint (Figure 1*a*). At comparable resolution, H atoms can be expected to be more visible in electron cryo-microscopy (cryo-EM) maps than in electron density maps due to the nature of the electrostatic potential (Clabbers & Abrahams, 2018; Maki-Yonekura *et al*., 2023). Yamashita *et al*. (2021) analysed H atom density from X-ray crystallographic and single particle analysis (SPA) cryo-EM data for apoferritin structures deposited in the PDB (Berman *et al*., 2000) and EMDB (Lawson *et al*., 2015) highlighting that even at 2.0 Å resolution it is possible to see some H atoms in cryo-EM maps. For extremely well-behaved samples, the recent ‘resolution revolution’ in SPA cryo-EM has allowed to achieve atomic resolution (Nakane *et al*., 2020; Yip *et al*., 2020). In the structure of apoferritin at 1.2 Å, most H atoms (approximately 70%) are easily discernible (Figure 1*b*). However, a recent microcrystal electron diffraction (microED) experiment on triclinic lysozyme reported at sub-atomic resolution, only allowed the identification of 35% of H atoms (Clabbers *et al*., 2022).

**Figure 1.**
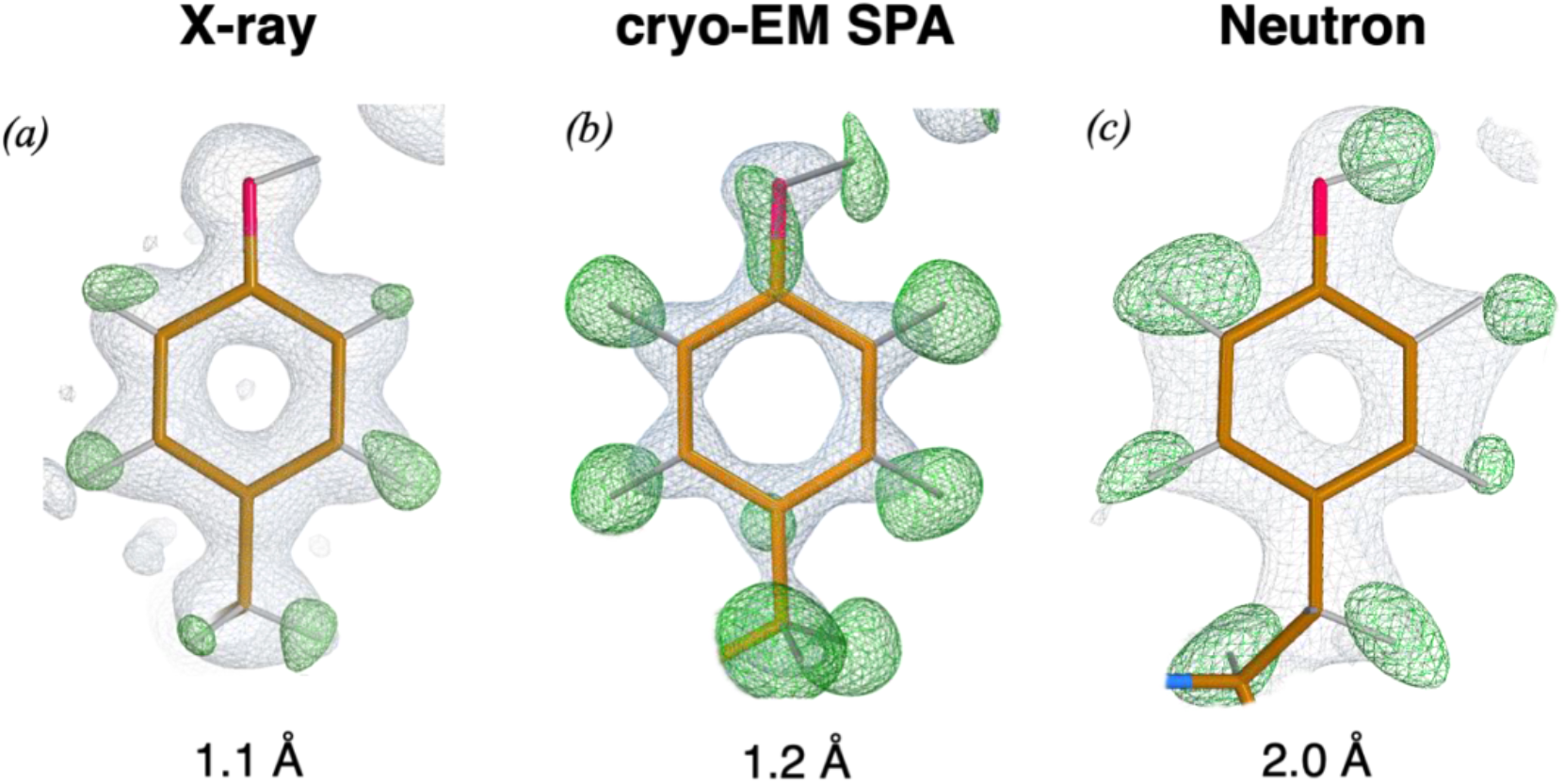
Map examples. *(a)* electron density maps for Tyr12 (PDB code: 3KYU, 1.1 Å resolution) show positive peaks for all aromatic H atoms, however, the H atom on the hydroxyl group is not visible; *(b)* cryo-EM maps for Tyr32 (PDB code: 7A4M, 1.22 Å resolution) show positive difference peaks for all H atoms; *(c)* neutron diffraction maps for Tyr146 (PDB code: 1CQ2, 2.0 Å resolution) show positive difference peaks for all H atoms (in the form of D), including the H atom on the hydroxyl group. Electron and neutron diffraction maps were calculated using *REFMAC5* (Murshudov *et al*., 2011), contoured at the +1.0*σ* (*2mF*_o_*−DF*_c_ in grey) and +3.0*σ* (*mF*_o_*−DF*_c_ in green) level. Cryo-EM weighted and sharpened *F*_o_ (grey) and omit (*F*_o_−*F*_c_ in green) maps were calculated using *Servalcat* (Yamashita *et al*., 2021) and contoured at the +1.5*σ* and +3.0*σ* level respectively. Molecular graphics representations were produced with *Coot 1.0* (Emsley *et al*., 2010).

Neutron macromolecular crystallography is a powerful technique that allows to directly visualise H atoms at more conventional resolution (Blakeley & Podjarny, 2018) Differently from X-rays that interact with atomic electron clouds, neutrons are scattered by nuclei (Fermi & Marshall, 1947). Atoms that are abundant in macromolecules typically possess positive coherent neutron scattering lengths (0.665×10^-12^ cm, 0.936×10^-12^ cm and 0.581×10^-12^ cm for C, N and O, respectively) that contribute favourably to the signal-to-noise (S/N) ratio of Bragg peaks. Although the scattering length of the common protium isotope (^1^H) is small and negative (-0.374×10^-12^ cm), its replacement with the heavier ^2^H isotope (D for deuterium, scattering length = 0.667×10^−12^ cm) makes them readily visible in neutron diffraction maps at 2.0-2.5 Å resolution or better (Figure 1*c*). Another important advantage of neutron diffraction for structure determination is the absence of global and specific radiation-induced damage that can be a serious limitation when using X-ray or electrons sources (Baker & Rubinstein, 2010; Garman, 2010).

Crystallographic refinement is one of the final step in the process of solving a macromolecular structure by diffraction methods (Tronrud, 2004). Various protocols are applied to maximise the agreement between diffraction data and model parameters that typically include atomic coordinates, atomic displacement parameters (ADPs), and occupancy values (Shabalin *et al*., 2018). Refinement of macromolecular models using neutron diffraction data can be currently carried out using packages initially developed for X-ray crystallographic refinement and modified to include neutron scattering lengths and the ability to deal with the refinement of individual H atom positions. They include the *nCNS* patch (Adams *et al*., 2009), an extension of the Crystallography and NMR System (*CNS*) package (Brünger *et al*., 1998) and *SHELXL2013* (Gruene *et al*., 2014). *SHELXL2013* is the most recent version of the *SHELXL* refinement program originally developed for small molecules and later adapted to macromolecules (Sheldrick, 2015). Another widely used package for neutron refinement is *phenix.refine* (Afonine *et al*., 2012) that is distributed as a part of the *PHENIX* suite (Adams *et al*., 2010). This program also includes the option of performing joint neutron/X-ray refinement, a concept introduced first in the field of small molecule crystallography (Coppens *et al*., 1981) and later applied to macromolecules with its *nCNS* implementation. Although effective joint neutron/X-ray refinement ideally requires the two datasets to be collected from the same crystal under the same conditions, it has the great advantage of increasing the available experimental data, thus compensating for the increased number of parameters arising from explicit addition of H atoms to the model.

Here, we describe an extension of the crystallographic refinement package *REFMAC*5 (Murshudov *et al*., 2011) of the *CCP*4 suite (Agirre *et al*., 2023) for the refinement of macromolecular models using neutron crystallographic data. Our implementation introduces a new parameter, dubbed “deuterium fraction”, representing the ^1^H/D fraction that is refined during the optimisation procedure. It also allows to effectively use stereochemical restraints from high-resolution reference structures, if available. We have tested *REFMAC*5 (version 5.8.0415) for the refinement of neutron models using ^1^H/D fraction parameters for selected or all H atoms together with restraints to a high-resolution known X-ray reference structure. Our evaluation involved the re-refinement of 97 PDB entries and one novel structure (FutA). The results of the refinement process are discussed in this study.

## 2. Methodology and Results

### 2.1. Reassessment of X–H restraint distances for macromolecular refinement

Macromolecular crystallographic refinement takes advantage of prior chemical knowledge. Information on ‘ideal’ bond lengths, bond angles and other chemical properties are incorporated into the target function and used in restrained refinement as subsidiary conditions to improve model parameters (Waser, 1963; Diamond, 1971; Jack & Levitt, 1978; Konnert & Hendrickson, 1980). Much of the available prior chemical knowledge used in macromolecular crystallographic refinement derives from high resolution small molecule X-ray diffraction experiments and the corresponding structures deposited in databases such as the Cambridge Structural Database (CSD) (Groom *et al*., 2016) or the Crystallography Open Database (COD) (Grazulis *et al*., 2012). For X–H (where X is a non-H ‘parent’ atom) bond lengths derived from X-ray diffraction experiments, their values reflect the relative positions of the atomic electron clouds. However, distances between H nuclei and their parent atoms are longer than those between electron clouds. This is because valence-electron density for H atoms is shifted toward their parent atoms (Coppens, 1997). Thus, to properly model and refine macromolecular models against neutron diffraction data, bond distance information should take this into account.

In addition to X-ray crystallographic structures, the CSD also contains a limited set of small molecule structures determined by neutron crystallography. Neutron entries in the CSD have almost doubled in recent years going from 1,213 in 2009 to 2,362 (1,452 organic and 910 metal-organic compounds) in 2021. An analysis of X–H bond lengths using the 2009 CSD neutron database was reported by Allen and Bruno (2010) that reassessed information derived from limited earlier data of the late 80s and early 90s (Allen *et al*., 1987; Orpen *et al*., 1989; Allen *et al*., 1992). We took advantage of the recent enrichment in neutron structures in the CSD and re-evaluated X–H bond length values. We employed the same approach of Allen and Bruno (2010) by selecting non-polymeric organic compounds without disorder and with *R-*factors ≤ 0.075 (647 entries). Entries derived from powder diffraction data were excluded. Neutron entries were retrieved using *ConQuest* (Bruno *et al*., 2002) and mean, median and standard deviation values for the X–H bond length distributions were estimated using *Mercury* (Macrae *et al*., 2020) O–H and N–H bond lengths were estimated by removing groups involved in very short hydrogen bonds as reported by Allen and Bruno (2010).

In an orthogonal approach we also derived X–H nuclear distances from quantum mechanics (QM) calculations. Initially, the stereochemical restraints generator *AceDRG* (Long *et al*., 2017) was employed to provide initial coordinates for 2,652 molecules constituted of twenty or fewer atoms selected from DrugBank (Wishart *et al*., 2018). The cut-off value on atom numbers was chosen to ensure computational efficiency while providing a pool size comparable to that of CSD entries. For geometry optimization, density functional theory (DFT) calculations were carried out with the self-consistent field wave function of restricted Hartree-Fock type as implemented in *GAMESS-US* (Schmidt *et al*., 1993). The hybrid generalised gradient approximation functional, B3LYP, was used with the (6-311++G**) basis set that includes both polarisation and diffuse functions. Solvent effect was calculated using the polarisable continuum model with water as solvent. More that 70% of the calculations ran successfully producing optimised coordinates for 1,874 molecules out of 2,652. We did not perform a detailed analysis of the calculations that ended prematurely.

Table 1 summarises nuclear bond distances for the most common X–H bond classes. It also provides values as reported by Allen and Bruno (2010) for reference. Overall, the recent nuclear bond distances derived from the CSD in 2021 are fully consistent with those previously derived in 2009. Nuclear distances obtained from theoretical calculations are also consistent with the experimentally derived values. The *AceDRG* data table has been updated to use median (*m*) and standard deviation (*σ*) values for all X–H nuclear distances from the CSD 2021 data (Figure 2*a*).

**Figure 2.**
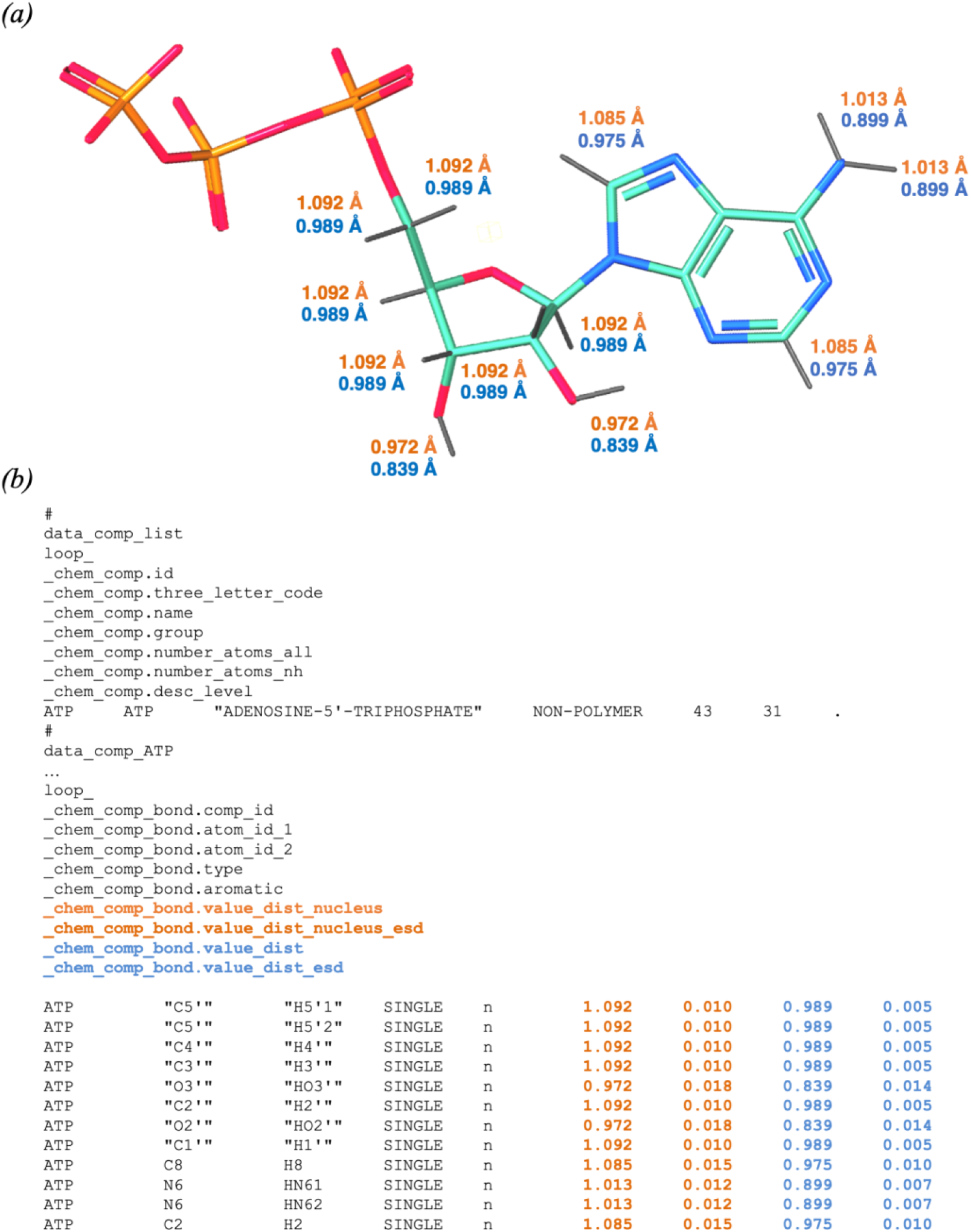
Example of a dictionary mmCIF file from the updated *AceDRG*. *(a)* 3D representation for the adenosine triphosphate (ATP) monomer. Nitrogen, carbon, oxygen, and phosphorus atoms are shown in blue, teal, red, and orange, respectively. X–H bonds are represented by grey sticks with their nuclear and X-ray diffraction-derived bond lengths (in Å) highlighted in orange and light blue, respectively; *(b)* Extract from the monomer description of the ATP component dictionary. The category _*chem_comp_bond* describes the bonded atoms, bond types and the ideal values of bond lengths and uncertainties associated with them. In this example, we show the ideal X–H bond lengths and standard deviations for nucleus positions (_*chem_comp_bond.value_dist_nucleus*, *chem_comp_bond.value_dist_nucleus_esd*) (orange) and electron positions (_*chem_comp_bond.value_dist*, _*chem_comp_bond.value_dist_esd*) (light blue).

**Table 1.**
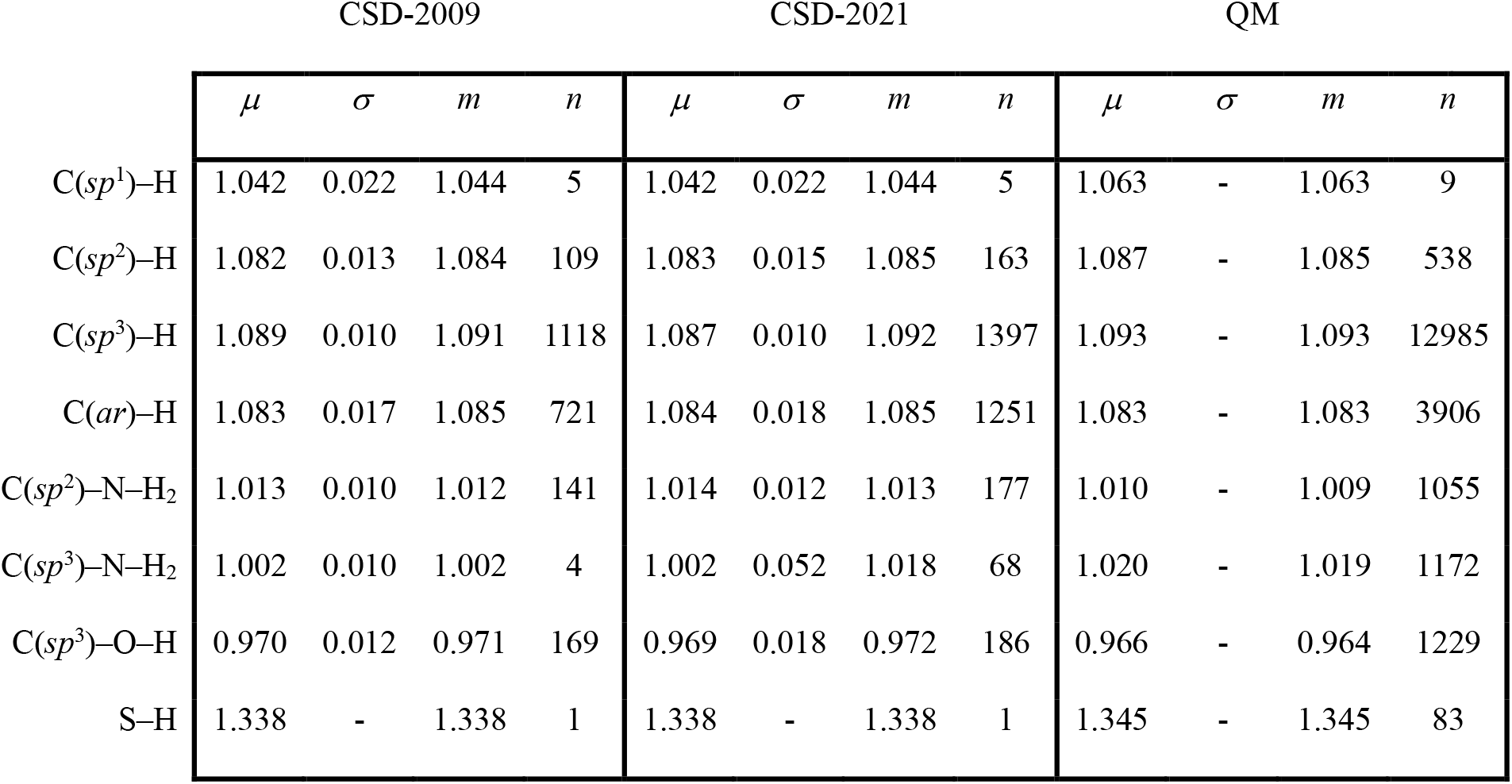
Nuclear X–H bond lengths. X represents (C, N, O, S) atoms covalently bound to H. For C, different types of hybridization are given with C(ar) indicating aromaticity. The first set of values (CSD-2009) is that of Allen and Bruno (2010) obtained from an analysis of curated neutron diffraction structures available within the CSD in 2009. The values in (CSD-2021) are updated values from our analysis using curated CSD entries as of 2021. The values in (QM) are the result of QM calculations carried out as described in the text. The letters μ, σ, m and n represent the mean, standard deviation, median, and number of observations for each X–H class, respectively. For QM, the standard deviation is not applicable. All bond lengths are in Å.

|  | CSD-2009 |  |  |  | CSD-2021 |  |  |  | QM |  |  |  |
| --- | --- | --- | --- | --- | --- | --- | --- | --- | --- | --- | --- | --- |
| | $\mu$ | $\sigma$ | $m$ | $n$ | $\mu$ | $\sigma$ | $m$ | $n$ | $\mu$ | $\sigma$ | $m$ | $n$ |
| C(*sp*<sup>1</sup>)–H | 1.042 | 0.022 | 1.044 | 5 | 1.042 | 0.022 | 1.044 | 5 | 1.063 | - | 1.063 | 9 |
| C(*sp*<sup>2</sup>)–H | 1.082 | 0.013 | 1.084 | 109 | 1.083 | 0.015 | 1.085 | 163 | 1.087 | - | 1.085 | 538 |
| C(*sp*<sup>3</sup>)–H | 1.089 | 0.010 | 1.091 | 1118 | 1.087 | 0.010 | 1.092 | 1397 | 1.093 | - | 1.093 | 12985 |
| C(ar)–H | 1.083 | 0.017 | 1.085 | 721 | 1.084 | 0.018 | 1.085 | 1251 | 1.083 | - | 1.083 | 3906 |
| C(*sp*<sup>2</sup>)–N–H<sub>2</sub> | 1.013 | 0.010 | 1.012 | 141 | 1.014 | 0.012 | 1.013 | 177 | 1.010 | - | 1.009 | 1055 |
| C(*sp*<sup>3</sup>)–N–H<sub>2</sub> | 1.002 | 0.010 | 1.002 | 4 | 1.002 | 0.052 | 1.018 | 68 | 1.020 | - | 1.019 | 1172 |
| C(*sp*<sup>3</sup>)–O–H | 0.970 | 0.012 | 0.971 | 169 | 0.969 | 0.018 | 0.972 | 186 | 0.966 | - | 0.964 | 1229 |
| S–H | 1.338 | - | 1.338 | 1 | 1.338 | - | 1.338 | 1 | 1.345 | - | 1.345 | 83 |

### 2.2. Inclusion of X–H nuclear distances in the *CCP*4 Monomer Library (*CCP*4-ML)

The *CCP*4-ML, also referred to as the *REFMAC*5 dictionary (Vagin *et al*., 2004), currently contains close to 35,300 entries for all standard and most non-standard amino acids, nucleotides, saccharides and various ligands. Each entry, identified as a monomer, possesses a unique code, and provides stereochemical information about constituent atoms, bonds distances, bond angles, torsion angles as well as stereochemical centres and planes. Statistics for these geometric parameters have been generated by *AceDRG* using data from COD. In addition, the *CCP*4-ML also contains more than 100 descriptors that specify covalent linkages between monomers and associated chemical modifications. The latter define all the chemical and geometric changes that occur to monomers following chemical reactions (for example removal of one of the oxygen atoms in peptide link formation). Covalent links refer to covalent interactions between monomers (for example, peptide links, sugar-peptide links, DNA/RNA links) (Nicholls *et al*., 2021a; Nicholls *et al*., 2021b).

The *CCP*4-ML has been recently updated to have X–H nuclear distances (orange in Figure 2*b*) as *_chem_comp_bond*.*value_dist_nucleus* and *value_dist_nucleus_esd* in addition to the distances between electron clouds (light blue in Figure 2*b*) (Nicholls *et al*., 2021b). X– H nuclear distances can now also be used to refine models from electron-derived experiments (SPA cryo-EM and microED), as both H atom ‘positions’ (electron and nucleus) contribute to the scattering.

### 2.3. *CCP*4 implementation of neutron macromolecular crystallographic refinement

#### 2.3.1. “Deuterium fraction” parametrisation

Neutron crystallographic experiments on macromolecules are typically carried out on ^1^H/D exchanged crystals to maximise the S/N ratio (Kossiakoff, 1984). This can be done by replacing exchangeable ^1^H atoms with D isotopes by soaking macromolecular crystals in deuterated media (Niimura & Podjarny, 2011). Alternatively, perdeuteration which replaces all ^1^H atoms with D isotopes can be carried out at the protein production stage by over-expressing protein(s) of interest in *Escherichia coli* or yeast strains in heavy water-based medium supplied with a perdeuterated carbon source, such as glycerol. Protein perdeuteration is a more effective method of improving the S/N as it dramatically lowers the incoherent background while enhancing the coherent scattering signal (Shu *et al*., 2000; Fisher *et al*., 2014). In addition, it avoids map cancellation issues due to protium’s negative scattering length (Blakeley & Podjarny, 2018; Logan, 2020). Currently, most neutron entries in the PDB (157 out of 213) reflect experiments carried out on partially deuterated samples as ^1^H/D exchange is simpler and less expensive than perdeuteration. However, the establishment of dedicated deuteration facilities and advanced experimental protocols have recently made perdeuteration more accessible to users (Meilleur *et al*., 2009; Budayova-Spano *et al*., 2020; Pierce *et al*., 2020).

In the refinement procedure implemented in *REFMAC*5, we have introduced a new quantity that represents the deuterium fraction for individual H atoms. This method is similar to the ‘deuterium saturation’ implemented in *SHELXL* (Gruene *et al*., 2014). In this parametrisation, protium ^1^H and deuterium D isotopes at each H position are not considered as separate entities. Instead, H atoms are represented by a unique set of coordinates that are associated with their isotope fraction which is optimised during the minimisation of the target function. The scattering factor for the ^1^H/D mixture is calculated using:

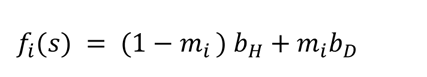

where *f*_*i*_(*S*) is the total contribution of protium and deuterium isotopes to the scattering factor of the *i*^-th^ H atom, *s* is the Fourier space vector, *b*_*H*_ is the deuterium fraction parameter that is an adjustable parameter, and *b*_*H*_ and *b*_*D*_ are the neutron scattering lengths of ^1^H and D isotopes, respectively. Neutron scattering lengths are tabulated in the *CCP*4 *atomsf_neutron* library, retrieved from https://www.ncnr.nist.gov/resources/n-lengths/list.html (Sears, 1992). The refined output model in mmCIF format contains only H atoms (no ^1^H/D or D sites) and a new *_atom_site.ccp4_deuterium_fraction* column representing the value of the deuterium fraction for each of the H atoms in the model. Users have the option to refine deuterium fraction parameters for either only polar or all H atoms. This method simplifies the model output as there is no ^1^H/D duplication for the same set of coordinates, for example, when alternative conformations are introduced in the structure (Figure 3*a*, *b*). The presence of only “generalised” H atoms with their corresponding deuterium fraction parameter, also reduces the risk of book-keeping errors. In the deuterium fraction representation, all D atoms are converted to H atoms and their presence is indicated by their corresponding deuterium fractions (Figure 3*c*, *d*). We note that this new item can be added only to mmCIF files, which is now the model deposition standard. For PDB files that have fixed column format, ^1^H and D are present at each H position and the deuterium fraction is indicated in the occupancy column.

**Figure 3.**
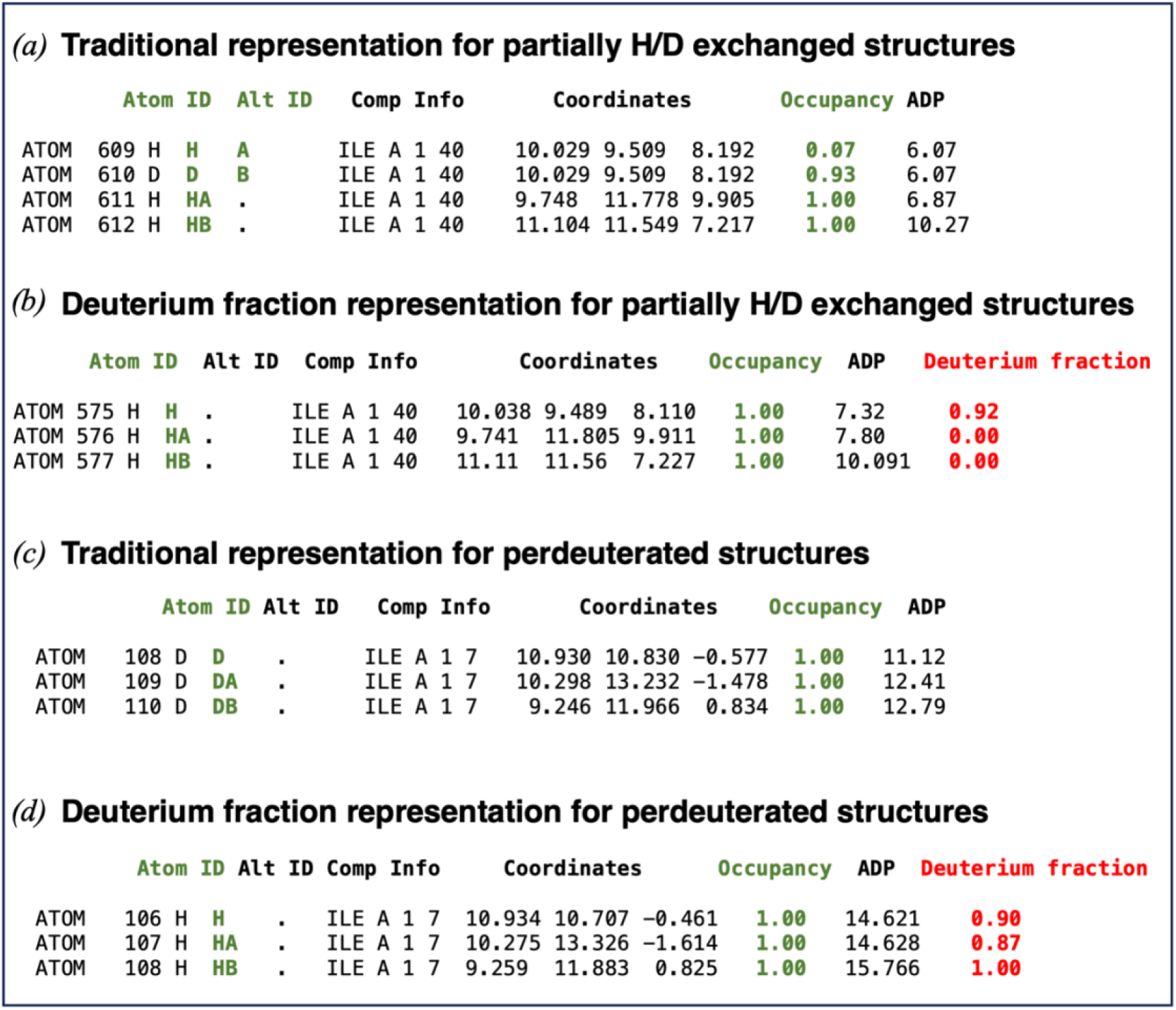
Comparison between the traditional representation for partially ^1^H/D exchanged structures and perdeuterated structures and the deuterium fraction representation in mmCIF files. *(a)* traditional exchangeable ^1^H/D sites representation extracted from the mmCIF file of 1VCX. The ^1^H atom bonded to the main-chain N atom of Ile40 is partially exchanged with D. ^1^H and D isotopes have a separate atom row in the atom table with alternative locations A and B (green). The sum of their total occupancy (green) is set to 1.0 (occupancy values of the ^1^H and D atoms are 0.07 and 0.93 respectively). The H atom bonded to the CA and CB of Ile40 are not exchanged during the partial deuteration procedure, their occupancy value is equal to 1.0. *(b)* deuterium fraction representation created by *REFMAC*5. A new column has been created that specifies the fraction of the deuterium substitution (where 100% is fully deuterated) for the H atoms exchanged. D atoms are not present in the atom table, only H atoms with the corresponding deuterium fraction parameters (red). The H atom bonded to the main-chain N atom of Ile40 has a deuterium fraction value of 0.92, while the H atom bonded to the CA and CB of Ile40 are not exchanged, hence the deuterium fraction for these H atoms is zero. *(c)* traditional perdeuterated sites representation extracted from the mmCIF file of 3RZ6. Here all the H atoms of Ile7 have been substituted with D atoms. There are no H atoms in the traditional perdeuterated structures. Occupancy of the D atoms is set to 1.0 by convention. *(d)* deuterium fraction representation for perdeuterated structures. All D atoms are converted to H atoms, the corresponding deuterium fractions are refined, and values close to 1.0 are obtained.

#### 2.3.2. Reference structure restraints

Neutron macromolecular crystallographic data often suffer from limited completeness and high-resolution is not always achievable. Therefore, a useful strategy to increase the data-to-parameter ratio in refinement is that of joint neutron/X-ray refinement provided that an isomorphous X-ray data set is available. This approach originally implemented in n*CNS* and available within *phenix.refine* of the *PHENIX* suite has been employed for the refinement of several macromolecular structures (Liebschner *et al*., 2018). Another possible strategy to increase the data-to-parameter ratio is the use of external restraints. This approach can be applied if a high-resolution model related to the target structure to be refined is available. With the use of external restraints, the target structure is pulled towards the conformation adopted by the known structure. Such restraints have been useful in the refinement of low-resolution X-ray (Headd *et al*., 2012; Nicholls *et al*., 2012; Smart *et al*., 2012; Schröder *et al*., 2014; Sheldrick, 2015; van Beusekom *et al*., 2018) and cryo-EM (Afonine *et al*., 2018; Nicholls *et al*., 2018) structures.

When performing neutron crystallographic studies of macromolecules, the corresponding high-resolution X-ray model is always available as invariably determined first. Thus, considering that X-ray models provide significantly more accurate coordinates for all non-H atoms, their use as source of external restraints appears as a useful approach to improve neutron refinement. The *CCP*4 program *ProSMART* (Nicholls *et al*., 2014) generates such external restraints, by distilling the local structure of a known reference model. Here, we used *ProSMART* to identify matching atoms by aligning the target model and an X-ray reference model, and then generated interatomic distance restraints between proximal non-H atoms within a given distance threshold (default 4.2 Å). The resulting external restraints were subsequently used by *REFMAC5* during the refinement of the target neutron model. We find that the injection of prior structural knowledge from homologous X-ray structures often results in more accurate and reliable neutron models.

### 2.4. Performance analysis by re-refinement of PDB entries

To test our current implementation, we re-refined 97 of the available neutron PDB entries (45.5% of the total) using *REFMAC*5. Of these, 55 are structures that were originally refined against neutron data only and 42 are entries deposited following a protocol of joint neutron/X-ray refinement. We selected our test pool based on the availability of experimental data (including complete cross-validation sets) and a wide resolution range (upper limit 1.05-2.75 Å).

For each entry, coordinate files (in PDB and mmCIF format) and crystallographic data (mmCIF format) were downloaded from the PDB. Each mmCIF reflection file was then converted into the MTZ format, which serves as the standard format used by *CCP*4 programs (Agirre *et al*., 2023). For the ‘neutron-only’ entries, the *CCP*4 program *CIF2MTZ* was utilised to convert mmCIF to MTZ format. For entries refined using a joint X-ray/neutron protocol, their mmCIF reflection files should contain two distinct data blocks - one for X-ray diffraction and one for neutron diffraction. However, a few entries have been erroneously deposited with a single dataset. Since the refinement process within *REFMAC5* was performed only against neutron reflections, those were extracted and converted to MTZ using *GEMMI* (Wojdyr, 2022). In case only intensities were available, they were converted into amplitudes using *Servalcat* (Yamashita *et al*., 2021) ’fw’ function, which implements the French-Wilson procedure (French & Wilson, 1978).

To compare refinement statistics with those reported in the PDB, all ^1^H and D atoms present in the models were retained without regeneration. *REFMAC5* is able to read ^1^H/D sites and D atoms using *Servalcat REFMAC5* controller (’refmacat’) that uses *GEMMI* for restraints generation (Yamashita *et al*., 2023). D atoms are converted to H atoms with the deuterium faction parameters by *GEMMI*, and their distances adjusted using nuclear values from the *CCP*4-ML. In cases such as 5KSC, where the original model does not contain any ^1^H (or D) atoms, except for water molecules, *GEMMI* was employed to add them at riding positions.

If H atoms are generated, it is necessary to initialise their deuterium fraction prior to refinement. Users can choose to initialise all H atoms or only polar ones. For perdeuterated structures, where all H atoms are replaced by D atoms, the initialisation process sets the deuterium fraction parameter to 1 for all H atoms. In the case of H/D-exchanged structures, the deuterium fraction is set to 1 only for H atoms exchanged with D. Subsequently, the refinement process is performed to optimise the deuterium fraction. Initialisation was not used for the refinement of most of the entries containing H/D or D sites, while it was necessary for few entries, such as 1C57, 1CQ2, 5KSC, and 1XQN, where only H atoms were present in the models.

Our standard refinement protocol consisted of five cycles of restrained positional and individual ADP refinement using the data in the published resolution range. Three cycles of deuterium fraction refinement were performed after each cycle of individual atomic refinement. For perdeuterated samples we allowed refinement of the deuterium fraction for all H atoms whilst for ^1^H/D-exchanged samples only polar H atoms had this parameter included in the optimisation. H atom positions have been refined individually with all available restraints (bond lengths, angles, planarity and torsion angles) to ensure proper geometry. We found that this procedure allows deuterium fraction parameters to converge as the models had been previously refined by the original depositors.

#### 2.4.1. Re-refinement of PDB entries originally refined against neutron data-only

Using the protocol described earlier, we re-refined with *REFMAC*5, 55 PDB entries that were originally refined using neutron data-only. Entries were chosen over a wide resolution range going from medium-low resolution (2.7 Å, PDB code 2EFA) to sub-atomic resolution (1.05 Å, PDB code 4AR3). *R*-factor statistics for all 55 re-refined models are given in Table 2.

**Table 2.** Re-refinement of selected neutron-only models from the PDB. Comparison of published *R*-factor statistics and those obtained by re-refinement using *REFMAC5*.

| PDB information |  |  | *REFMAC5* refinement statistics |  |  |  |
| --- | --- | --- | --- | --- | --- | --- |
| PDB code | Published resolution range (low-high) (Å) | Published *R* values (work/free) (%) | Initial *R* values (work/free) (%) | Final *R* values (work/free) (%) | Data completeness (%) | No. of reflections |
| 1C57 | 15.79-2.40 | 27.0/30.1 | 29.7/33.0 | 19.9/25.4 | 87.38 | 8129 |
| 1CQ2 | 6.00-2.00 | 16.0/25.0 | 18.6/25.7 | 14.9/24.7 | 91.07 | 7528 |
| 1IU6 | 10.00-1.60 | 20.1/22.8 | 21.1/23.4 | 18.4/22.9 | 87.16 | 5775 |
| 1V9G | 25.05-1.80 | 22.2/29.4 | 26.9/33.5 | 22.9/33.7 | 85.31 | 1949 |
| 1VCX | 27.60-1.50 | 18.6/21.7 | 19.0/21.7 | 17.8/21.8 | 81.94 | 6620 |
| 1WQ2 | 20.00-2.40 | 22.9/28.9 | 28.6/32.1 | 21.5/28.7 | 92.29 | 6232 |
| 1XQN | 32.82-2.50 | 26.6/32.0 | 30.0/31.5 | 23.0/29.4 | 74.99 | 6088 |
| 2DXM | 8.00-2.10 | 19.7/26.0 | 20.2/26.6 | 18.8/26.4 | 64.12 | 20178 |
| 2EFA | 80.00-2.70 | 21.6/29.1 | 24.3/27.6 | 22.3/28.0 | 95.66 | 2154 |
| 2GVE | 10.00-2.20 | 26.8/31.9 | 26.3/30.3 | 23.4/29.8 | 93.73 | 22133 |
| 2WYX | 19.41-2.10 | 22.3/25.8 | 25.8/28.7 | 22.5/27.7 | 86.85 | 15033 |
| 2XQZ | 53.19-2.10 | 22.5/25.9 | 25.9/29.1 | 19.0/26.5 | 77.26 | 13374 |
| 2YZ4 | 33.64-2.20 | 27.9/31.2 | 28.1/31.3 | 22.0/27.6 | 66.06 | 8044 |
| 2ZOI | 70.00-1.50 | 19.2/21.9 | 19.9/22.4 | 18.5/21.5 | 89.64 | 14526 |
| 2ZPP | 20.00-2.50 | 22.1/26.0 | 22.8/26.8 | 20.5/25.9 | 98.61 | 2612 |
| 2ZWB | 20.00-1.80 | 22.3/24.7 | 22.8/25.0 | 18.0/23.1 | 95.93 | 9862 |
| 3A1R | 30.84-1.70 | 19.5/23.8 | 18.5/22.9 | 16.1/21.8 | 81.89 | 10300 |
| 3FHP | 41.43-2.00 | 16.8/24.7 | 18.6/23.4 | 16.8/22.2 | 81.46 | 4615 |
| 3KMF | 20.00-2.00 | 25.0/30.0 | 28.9/29.1 | 27.9/31.0 | 86.35 | 31611 |
| 3Q3L | 36.42-2.50 | 22.1/26.8 | 23.0/26.8 | 22.9/26.6 | 73.25 | 27157 |
| 3RYG | 27.11-1.75 | 18.1/20.0 | 22.8/23.8 | 20.4/25.5 | 92.21 | 4884 |
| 3RZ6 | 21.77-1.75 | 20.8/23.8 | 25.2/25.6 | 21.3/25.1 | 79.55 | 4215 |
| 3RZT | 27.21-1.75 | 20.2/24.9 | 25.4/29.3 | 21.9/29.2 | 75.6 | 3989 |
| 3SS2 | 24.33-1.75 | 21.1/24.2 | 23.9/26.2 | 20.1/26.0 | 77.19 | 4080 |
| 3U2J | 12.10-2.00 | 23.2/27.2 | 23.5/27.0 | 22.4/26.5 | 86.54 | 14521 |
| 4AR3 | 15.71-1.05 | 19.9/23.7 | 19.7/23.0 | 18.8/22.4 | 88.73 | 21580 |
| 4AR4 | 27.46-1.38 | 18.6/22.6 | 17.9/22.1 | 15.5/21.0 | 91.56 | 9958 |
| 4BD1 | 9.97-2.00 | 21.9/25.7 | 22.4/25.0 | 14.6/17.6 | 87.72 | 18290 |
| 4C3Q | 10.00-2.20 | 19.2/24.0 | 21.3/25.1 | 17.3/24.2 | 88.19 | 13256 |
| 4FC1 | 10.00-1.10 | 21.1/25.3 | 22.8/25.5 | 21.5/24.1 | 76.85 | 10549 |
| 4G0C | 20.00-2.00 | 26.7/28.3 | 27.2/26.0 | 24.3/28.1 | 84.4 | 13742 |
| 4K9F | 27.14-1.75 | 19.9/24.1 | 21.6/26.0 | 17.8/25.7 | 74.37 | 3671 |
| 4Q49 | 20.00-1.80 | 18.7/21.5 | 18.6/20.9 | 15.5/20.4 | 89.45 | 18803 |
| 4Y0J | 20.00-2.00 | 26.3/29.1 | 29.2/30.6 | 26.6/31.8 | 80.89 | 13095 |
| 4ZZ4 | 51.19-1.80 | 19.7/22.1 | 19.5/20.6 | 17.7/23.4 | 71.33 | 7524 |
| 5A90 | 38.91-1.70 | 19.2/22.7 | 21.1/24.1 | 18.9/23.2 | 88.61 | 28772 |
| 5GX9 | 33.45-1.49 | 15.8/20.0 | 17.3/21.0 | 17.0/20.9 | 93.5 | 14375 |
| 5KSC | 10.00-2.10 | 24.0/28.0 | 29.6/31.4 | 23.5/30.9 | 70.78 | 12424 |
| 5MNX | 22.17-1.42 | 16.6/20.6 | 18.5/21.9 | 18.3/21.8 | 90.57 | 36197 |
| 5MNY | 22.09-1.43 | 16.4/19.3 | 18.1/20.5 | 18.2/20.2 | 93.83 | 36397 |
| 5MNZ | 21.32-1.45 | 16.9/20.1 | 18.1/21.1 | 18.0/21.0 | 90.2 | 33722 |
| 5MO1 | 19.68-1.49 | 17.5/21.6 | 18.6/22.3 | 18.9/22.5 | 93.38 | 32293 |
| 5MO2 | 22.14-1.50 | 16.1/20.0 | 17.4/20.8 | 17.2/20.5 | 88.77 | 30027 |
| 5TY5 | 14.78-2.30 | 23.9/25.2 | 27.2/27.0 | 24.4/29.8 | 73.88 | 34761 |
| 5VG1 | 12.00-2.10 | 18.7/26.5 | 21.4/27.5 | 19.8/27.0 | 75.47 | 13304 |
| 5VNQ | 16.70-2.20 | 24.2/28.0 | 27.5/30.1 | 23.7/29.0 | 71.29 | 7734 |
| 5ZO0 | 22.86-1.65 | 18.6/22.9 | 19.7/23.2 | 18.2/22.4 | 86.21 | 21422 |
| 6C78 | 14.76-1.75 | NA/NA | 23.9/25.6 | 19.8/24.9 | 85.31 | 25773 |
| 6GTJ | 32.66-1.80 | 23.2/27.6 | 24.7/28.6 | 19.7/27.6 | 78.58 | 21858 |
| 6H1M | 21.76-2.15 | 21.8/24.9 | 22.3/25.3 | 18.5/22.5 | 91.42 | 12707 |
| 6L26 | 24.76-1.44 | 16.8/20.6 | 18.9/22.1 | 16.9/21.0 | 88.73 | 33763 |
| 7JOR | 17.23-2.05 | 24.9/28.8 | 25.4/29.5 | 23.2/29.5 | 79.44 | 12714 |
| 7KKS | 14.64-2.20 | 25.7/28.2 | 28.1/30.7 | 22.4/29.0 | 98.34 | 23326 |
| 7KKW | 14.65-2.30 | 24.9/30.2 | 27.9/31.8 | 22.5/31.6 | 98.38 | 20628 |
| 7VEI | 17.70-2.00 | 17.0/21.5 | 16.8/21.5 | 14.2/21.3 | 98.3 | 7555 |

For some entries (e.g., 1WQ2, 3RZ6, 4C3Q, 4FC1, 5A90, 5GX9, 7KKW in Table 2), we observe that initial *R*_work_ and *R*_free_ values are higher than those reported in the PDB. In the case of 1WQ2, the PDB header reports 22.9% and 28.9% for *R*_work_ and *R*_free_, respectively, while the paper indicates values of 28.2 and 30.1% (Chatake *et al*., 2003). The latter values are similar to the initial *R*-factors from *REFMAC5* (28.6% and 32.1% for *R*_work_ and *R*_free_, respectively). Following refinement, *R*_work_ and *R*_free_ values from *REFMAC5* become comparable to the deposited values suggesting convergence of the refinement procedure (Table 2).

For several structures including the low resolution 1C57, 1WQ2, 1XQN, 2GVE, 2YZ4 entries, medium resolution 3FHP, 3U2J, 4BD1, 6H1M entries and high resolution 2ZOI, 2ZWB, 3A1R, 4AR3, 4AR4, 4FC1, 4Q49 entries, *R*_work_/*R*_free_ values obtained from *REFMAC5* are lower compared to the deposited ones, improving often by ∼2–3 percentage points. However, for a few other entries, the final *R*-factor values obtained from *REFMAC5* are slightly higher. One explanation is that, in this study, the models have been re-refined without any additional refinement strategy that could significantly improve the refinement statistics. For example, the application of TLS refinement (Winn *et al*., 2001; Winn *et al*., 2003), as well as the use of anisotropic ADP refinement for high resolution structures and jelly-body restraints, could potentially improve the refined model. One general point of consideration, however, is that the calculation of scaling factors used in the *R*-factor equation is different among refinement packages and this can lead to differences in *R*-factors. Although overall *R*-values are not the only metrics to consider when evaluating the quality of a structural model which cannot be properly assessed without careful map analysis, the values obtained from this test set indicate that our implementation for the neutron crystallographic refinement performs satisfactorily.

#### 2.4.2. Re-refinement of PDB entries originally refined using a joint neutron/X-ray

We also tested refinement of 42 models previously obtained through joint neutron/X-ray refinement utilizing solely neutron data and incorporating the deuterium fraction parameterisation. Table 3 presents all joint neutron/X-ray models, featuring neutron data from lowest to highest resolutions, selected for re-refinement within *REFMAC5*. The table compares the *R*-factor statistics published for these selected entries with the *R*-factors obtained through their re-refinement using *REFMAC5*.

**Table 3.** Re-refinement of selected joint neutron/X-ray models from the PDB using neutron diffraction data only. Comparison of published *R*-factor statistics and those obtained by re-refinement using *REFMAC5*.

| PDB information |  |  | *REFMAC5* refinement statistics |  |  |  |
| --- | --- | --- | --- | --- | --- | --- |
| PDB code | Published resolution range (low-high) (Å) | Published *R* values (work/free) (%) | Initial *R* values (work/free) (%) | Final *R* values (work/free) (%) | Data completeness (%) | No. of reflections |
| 2R24 | 40.11-2.19 | 25.7/29.1 | 25.9/29.7 | 22.1/30.0 | 72.76 | 10892 |
| 3R98 | 53.54-2.40 | 20.7/25.1 | 20.8/26.1 | 18.4/25.7 | 74.99 | 11610 |
| 3R99 | 53.54-2.40 | 20.7/25.0 | 20.8/26.1 | 18.5/25.6 | 74.99 | 11610 |
| 3VXF | 44.87-2.75 | 18.3/23.4 | 17.7/23.6 | 17.0/24.3 | 73.72 | 7670 |
| 3X2O | 18.83-1.50 | 22.8/25.1 | 23.6/25.6 | 21.1/24.0 | 93.49 | 23109 |
| 3X2P | 19.68-1.52 | 21.8/26.0 | 22.8/26.6 | 22.0/25.5 | 91.46 | 25101 |
| 4CVI | 39.84-2.41 | 17.6/24.3 | 17.6/23.8 | 14.8/22.9 | 73.74 | 11426 |
| 4DVO | 20.00-2.00 | 19.0/21.4 | 19.8/22.1 | 17.9/22.6 | 91.62 | 28663 |
| 4GPG | 37.61-1.98 | 19.5/25.9 | 20.0/25.9 | 20.0/26.0 | 69.97 | 10446 |
| 4NY6 | 26.77-1.85 | 17.6/22.4 | 19.6/22.3 | 15.2/21.1 | 89.97 | 4654 |
| 4PDJ | 32.40-1.99 | 23.0/27.1 | 23.6/25.9 | 20.4/26.1 | 78.41 | 8315 |
| 4PVM | 36.38-2.00 | 20.9/27.1 | 21.4/27.1 | 19.0/27.4 | 76.96 | 12062 |
| 4PVN | 52.28-2.30 | 20.9/26.2 | 21.4/26.8 | 18.6/27.0 | 98.62 | 10831 |
| 4QCD | 21.19-1.93 | 16.7/22.7 | 17.8/22.5 | 17.5/22.6 | 79.33 | 16540 |
| 4QDW | 20.00-1.80 | 16.6/17.9 | 19.2/18.2 | 16.2/17.5 | 72.86 | 31459 |
| 4S2G | 20.00-2.00 | 16.4/18.2 | 17.5/18.4 | 14.4/18.8 | 93.51 | 13125 |
| 4XPV | 20.00-2.00 | 26.4/30.4 | 27.3/30.0 | 24.8/29.9 | 80.58 | 11251 |
| 5A93 | 15.03-2.20 | 21.7/23.6 | 23.0/22.8 | 15.9/23.7 | 71.94 | 11004 |
| 5CG5 | 53.57-2.40 | 18.6/22.9 | 19.3/22.9 | 17.5/22.8 | 98.45 | 17458 |
| 5CG6 | 22.12-2.40 | 26.0/28.7 | 26.0/25.3 | 24.2/27.0 | 98.25 | 17245 |
| 5JPR | 36.71-2.20 | 23.6/31.0 | 23.6/31.0 | 20.2/31.2 | 68.13 | 8761 |
| 5MON | 22.13-1.42 | 17.0/18.1 | 20.7/21.4 | 18.2/20.9 | 90.56 | 36196 |
| 5MOO | 22.09-1.43 | 17.0/18.5 | 19.9/20.9 | 18.0/20.3 | 93.83 | 36397 |
| 5MOP | 21.33-1.45 | 17.2/18.4 | 19.4/20.6 | 17.7/20.7 | 90.2 | 33722 |
| 5MOQ | 25.43-1.50 | 15.0/16.7 | 17.1/18.5 | 15.3/18.5 | 89.39 | 30123 |
| 5MOR | 19.67-1.49 | 19.6/20.7 | 22.8/23.8 | 18.9/22.0 | 93.38 | 32293 |
| 5MOS | 22.15-1.50 | 16.6/18.0 | 18.7/19.5 | 16.9/19.8 | 88.77 | 30027 |
| 5XPE | 17.02-2.09 | 22.5/27.8 | 24.2/28.2 | 21.0/28.7 | 78.13 | 9595 |
| 5ZN0 | 33.76-1.90 | 18.8/24.7 | 19.6/24.9 | 17.4/25.6 | 97.05 | 22817 |
| 6BBR | 15.00-2.30 | 22.1/24.6 | 25.8/25.1 | 21.4/26.1 | 90.97 | 7336 |
| 6BBZ | 15.43-2.20 | 19.8/22.0 | 24.2/24.0 | 19.7/24.6 | 92.18 | 8108 |
| 6BQ8 | 40.00-2.20 | 23.2/28.8 | 22.8/28.4 | 20.5/30.6 | 72.77 | 5686 |
| 6EXY | 28.20-1.70 | 15.0/18.7 | 18.6/19.5 | 15.8/19.5 | 95.37 | 14401 |
| 6U0C | 15.00-2.10 | 21.8/25.0 | 23.2/27.1 | 19.0/26.3 | 84.10 | 10060 |
| 6U0E | 14.51-1.89 | 21.7/25.4 | 26.1/28.0 | 21.8/26.9 | 91.19 | 14921 |
| 6U0F | 13.79-2.00 | 21.9/24.1 | 25.2/26.5 | 20.6/25.9 | 86.60 | 12087 |
| 7A0L | 40.00-2.10 | 21.5/23.0 | 23.0/23.2 | 17.0/26.0 | 78.62 | 18131 |
| 7D6G | 26.33-2.10 | 17.7/21.9 | 20.3/22.1 | 16.3/22.4 | 71.99 | 6598 |
| 7JUN | 14.96-2.50 | 20.1/25.3 | 20.3/25.2 | 17.9/26.5 | 83.14 | 7617 |
| 7TX3 | 14.12-1.89 | 22.6/27.5 | 23.0/27.9 | 21.8/28.5 | 93.70 | 22968 |
| 7TX4 | 12.75-2.35 | 17.7/25.9 | 17.9/25.9 | 16.4/26.1 | 85.83 | 5371 |
| 7TX5 | 32.87-2.30 | 18.7/25.9 | 19.4/26.5 | 18.1/27.1 | 75.39 | 5882 |

The *R*_work_ and *R*_free_ values obtained from *REFMAC*5 (Table 3, Final *R* values work/free column) by refining joint models using only neutron data were found to be similar to those obtained from joint neutron/X-ray refinement (Table 3, Published *R* values work/free column). For some entries, the *R*-factors are slightly improved compared to the published ones. It is widely acknowledged that refinement solely using neutron data may lead to overfitting due to the explicit refinement of H atom parameters. However, the gap observed between the *R*_work_ and *R*_free_ values obtained from *REFMAC*5 is not substantial (mean Δ*R* is 5%). Thus, this strategy can be a viable alternative when joint refinement is not feasible.

#### 2.4.3. Re-refinement using external restraints

To improve the quality of neutron atomic models, especially at low resolution, a re-refinement has been performed by incorporating X-ray reference structure restraints. A subset of models obtained by neutron refinement only and from joint neutron/X-ray, featuring neutron data at low resolution and few at high resolution were selected for this analysis. For the ‘neutron-only’ entries (1C57, 2EFA, 2YZ4 and 2ZPP), the corresponding X-ray reference structures were chosen from the PDB based on their high structural similarity to the neutron refined structures. The ‘Find Similar Assemblies’ option from the PDB uses the Structure Similarity Search (Guzenko *et al*., 2020) to assess global 3D-shape similarity providing a Structure Match Score indicating the probability in percentage that the structure match is similar to the query. The X-ray structures chosen reported the highest Structure Match Score. If a suitable X-ray reference model is not known, we recommend running a BLAST (Altschul *et al*., 1997) search over the whole PDB by inputting a FASTA sequence of the target neutron model.

For the joint neutron/X-ray structures selected a different protocol was applied. Firstly, these models were subjected to refinement against their corresponding X-ray data using *REFMAC5*, with a total of ten refinement cycles. The output model obtained from this refinement process was subsequently employed as a reference model.

*ProSMART* (Nicholls *et al*., 2014) takes as input the neutron target model and X-ray reference structure model in PDB or mmCIF format and generates interatomic distance restraints between proximal non-H atoms reported in a restraint file. The refinement was performed by simultaneously refining non-H atoms of the model by using restraints generated by *ProSMART* and by using the deuterium fraction parametrisation for H atoms (twenty refinement cycles interleaved with three deuterium fraction refinement^1^). PDB information for the neutron and X-ray models selected, as well as published refinement statistics and those obtained by *REFMAC5* are shown in Table 4.

**Table 4.** Selected neutron models and corresponding X-ray reference models for re-refinement within *REFMAC5* using external restraints. Comparison of published *R*-factor statistics and those obtained by re-refinement using *REFMAC5*.

| PDB code |  | Published resolution range (low-high) (Å) |  | Published *R* values (work/free) (%) |  | *REFMAC5* *R* values (work/free) (%) |
| --- | --- | --- | --- | --- | --- | --- |
| Neutron | X-ray | Neutron | X-ray | Neutron | X-ray | Neutron refinement with external restraints |
| 1C57 | 1DQ6 | 15.79-2.40 | 8.00-1.90 | 27.0/30.1 | 18.6/NA | 20.2/24.9 |
| 2EFA | 1B2A | 80.00-2.50 | 55.00-1.70 | 21.6/29.1 | 18.8/23.0 | 21.9/25.3 |
| 2YZ4 | 1DQ6 | 33.64-2.20 | 8.00-1.90 | 27.9/31.2 | 18.6/NA | 21.7/27.3 |
| 2ZPP | 1B2G | 20.00-2.50 | 10.00-1.80 | 22.1/26.0 | 20.0/22.6 | 20.8/25.4 |
| 3R98 | 3R98 | 53.54-2.40 | 43.89-2.10 | 20.7/25.1 | 16.6/20.3 | 17.8/25.2 |
| 3R99 | 3R99 | 53.54-2.40 | 43.89-2.10 | 20.7/25.0 | 16.6/20.3 | 17.9/25.3 |
| 3VXF | 3VXF | 44.87-2.75 | 29.22-1.60 | 18.3/23.4 | 16.1/18.4 | 16.8/23.5 |
| 3X2O | 3X2O | 18.83-1.50 | 28.23-1.00 | 22.8/25.1 | 13.5/15.3 | 21.2/23.9 |
| 3X2P | 3X2P | 19.68-1.52 | 37.75-0.99 | 21.8/26.0 | 13.4/14.2 | 22.0/25.7 |
| 4CVI | 4CVI | 39.84-2.41 | 14.80-2.10 | 17.6/24.3 | 13.4/17.6 | 14.0/22.8 |
| 4DVO | 4DVO | 20.00-2.00 | 29.90-1.55 | 19.0/21.4 | NA/NA | 18.4/22.2 |
| 4GPG | 4GPG | 37.61-1.98 | 50.00-1.89 | 19.5/25.9 | 14.6/20.3 | 20.9/25.7 |
| 4PVN | 4PVN | 52.28-2.30 | 43.13-1.95 | 20.9/26.2 | 15.6/18.5 | 18.7/26.8 |
| 5CG6 | 5CG6 | 22.12-2.40 | 44.31-1.70 | 26.0/28.7 | 19.6/21.1 | 24.0/27.8 |
| 5XPE | 5XPE | 17.02-2.09 | 46.50-1.64 | 22.5/27.8 | 15.5/18.5 | 20.5/29.7 |
| 5ZN0 | 5ZN0 | 33.76-1.90 | 36.01-1.10 | 18.8/24.7 | 18.6/21.2 | 16.7/25.5 |
| 6BQ8 | 6BQ8 | 40.00-2.20 | 10.00-2.00 | 23.2/28.8 | 19.9/24.5 | 19.7/29.2 |
| 6EXY | 6EXY | 28.20-1.70 | 31.93-1.10 | 15.0/18.7 | 12.3/13.7 | 16.0/19.8 |
| 6U0E | 6U0E | 14.51-1.89 | 29.01-2.10 | 21.7/25.4 | 18.4/23.5 | 21.4/26.7 |
| 7D6G | 7D6G | 26.33-2.10 | 40.00-1.65 | 17.7/21.9 | 15.7/18.6 | 14.6/22.1 |
| 7TX4 | 7TX4 | 12.75-2.35 | 61.05-1.90 | 17.7/25.9 | 16.6/22.4 | 15.2/26.0 |

The incorporation of external restraints has been observed to improve both the *R*_work_ and *R*_free_ values for low resolution neutron structures. Specifically, *R*-factors are improved by ∼2–3 percentage points in certain cases (1C57, 2EFA, 2YZ4, 4CVI) (Table 4, Neutron refinement with external restraints) compared with both the published values and those obtained using deuterium fraction refinement only. Moreover, certain high-resolution structures (3X2O, 3X2P) also demonstrate improved *R*-factors, which indicate that these restraints can improve the quality of neutron models regardless of the resolution.

### 2.5. Selected examples of neutron crystallographic refinement

#### 2.5.1. Re-refinement of the neutron structure of chloride-free urate oxidase (UOX) in complex with its 8-azaxanthine (8AZA) inhibitor

Our re-refinement runs reported in previous paragraphs looked mainly at global refinement statistics. As a selected example that involved a more detailed inspection of neutron maps we carried out the re-refinement of the joint neutron/X-ray structure of perdeuterated urate oxidase (UOX) in complex with its 8-azaxanthine (8AZA) inhibitor (PDB entry 7A0L) (McGregor *et al*., 2021).

In many organisms, the degradation of uric acid (UA) to 5-hydroxyisourate (5-HIU) is catalysed by cofactor-independent UOX (Kahn *et al*., 1997). In a two-step reaction, first, UA reacts with O_2_ to yield dehydroisourate (DHU) via a 5-peroxoisourate intermediate (Bui *et al*., 2014). This is then followed by a hydration step, in which DHU is hydroxylated to 5-HIU (Kahn, 1999; Wei *et al*., 2017).The joint structure of perdeuterated UOX in complex with its 8AZA inhibitor, relevant to the hydration step, has been recently determined using X-ray and neutron data at 1.33 and 2.10 Å resolution, respectively (McGregor *et al*., 2021). Joint refinement was carried out with *phenix.refine* (Afonine *et al*., 2010). It showed that the catalytic water molecule (W1) is present in the peroxo hole as neutral H_2_O (D_2_O), oriented at 45° with respect to the organic ligand. It is stabilized by Thr57 and Asn254 on different UOX protomers as well as by an O–H×××p interaction with 8AZA. The active site Lys10–Thr57 dyad features a charged Lys10–NH ^+^ side chain engaged in a strong hydrogen bond with Thr57^OG1^, while the Thr57^OG1–HG1^ bond is oriented toward the π system of the ligand, on average.

Re-refinement of the UOX:8AZA complex with *REFMAC5* was performed against neutron data alone using deuterium fraction parameterisation and external restraints. H and D atoms on residues and water molecules previously modelled were maintained at their positions and not regenerated. Deuterium fraction parameters were refined for all H atoms. H atom positions were refined individually. External restraints were generated using *ProSMART*, by firstly, re-refining the model against its X-ray data (10 cycles) and using the output model as a reference structure. Data collection and refinement statistics are given in Table S1.

8AZA is bound as a monoanion deprotonated at N3 and omit neutron maps confirm that W1 is neutral (Figure 4*a*). This supported by the presence of positive peaks for two deuterium atoms whose deuterium fraction values refine at 0.77 (H1) and 0.84 (H2). The protonation state of the Lys10–Thr57 active-site dyad has also been investigated. Omit neutron maps for Lys10 show that the residue is positively charged due to the presence of a ‘tri-lobe’ density distribution around NZ (Figure 4*b*). All hydrogen atoms bound to Lys10 refine with a high deuterium fraction parameter value (> 0.80). The direction of the OG1-HG1 bond in Thr57 was not easily identified in the original work (McGregor *et al*., 2021). Here, omit maps reveal positive density for Thr57^HG1^ at the 2.5*σ* level (Figure 4*b*). We refined the orientation of the OG1-HG1 bond by using the *REFMAC5* ‘hydrogen refine rpolar’ (rotatable polar) option, resulting in an optimal fit to the density. The deuterium fraction parameter for HG1 refined at 0.81. The orientation of the OG1-HG1 bond suggests the formation of another O–H×××p interaction with the N7 of 8AZA at 2.56 Å and a hydrogen bond is also formed between Lys10^HZ1^ and Thr57^OG1^ at a distance of 1.87 Å (Figure 4*b*). Overall, our results are fully consistent with those from the previous study (McGregor *et al*., 2021) and mechanistic considerations can be found therein.

**Figure 4.**
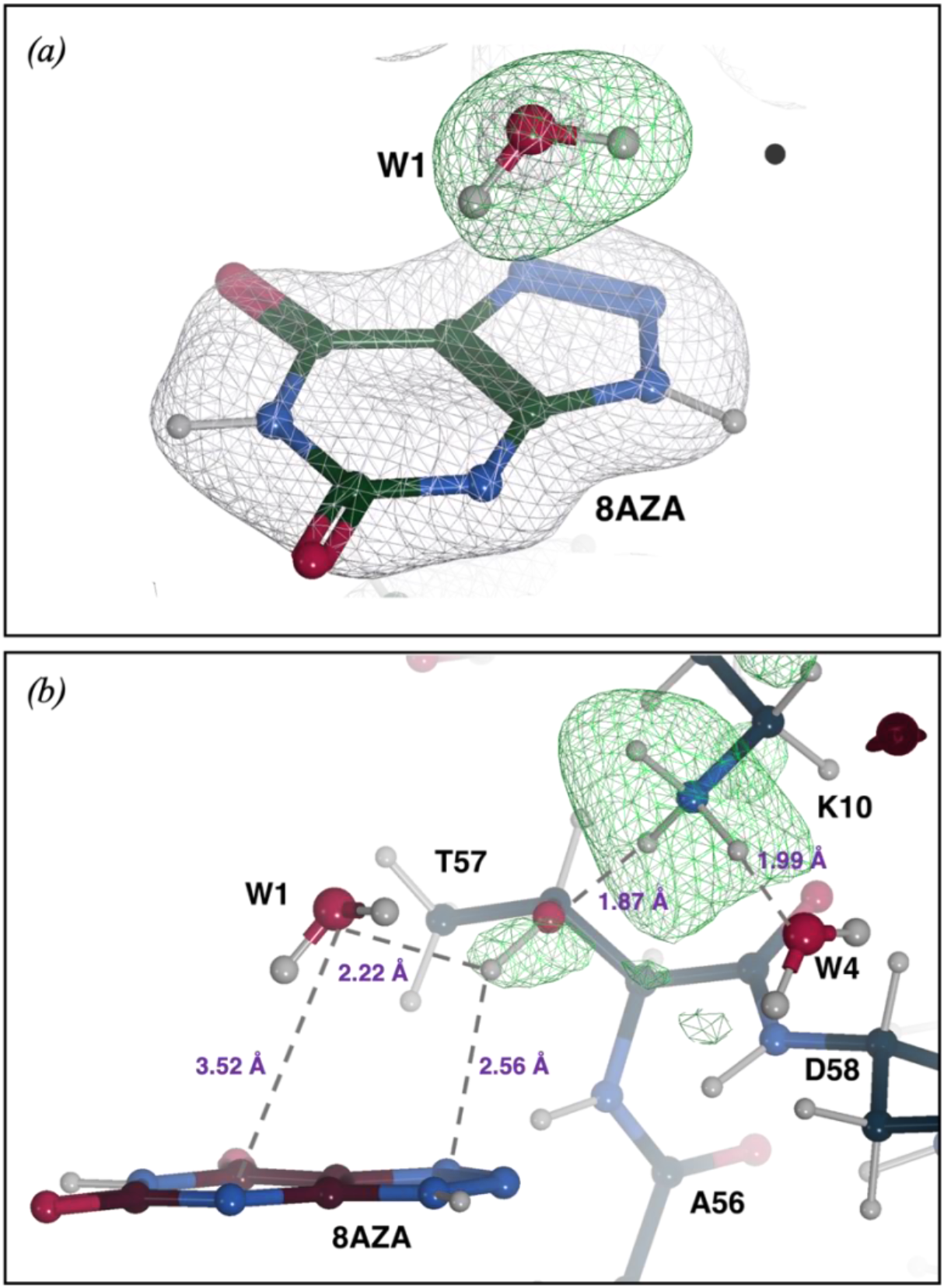
Neutron structure of the UOX-8AZA complex. *(a)* W1 is present as a neutral H_2_O molecule. *2mF*_o_*−DF*_c_ neutron diffraction map for 8AZA and the oxygen atom of W1 is shown in grey at the +1.0*σ* level. An omit *mF*_o_*−DF*_c_ neutron map indicates the presence of two deuterons (H1, H2) as suggested by the elongated positive density (in green at the +3.0*σ* level) next to the oxygen atom. *(b)* representation of a portion of the active site highlighting the protonation of Lys10–Thr57 dyad and the H-bond network. H-difference neutron density for Lys10^NZ^ and Thr57^OG1^ is shown in green at the +3.0*σ* and +2.5*σ* levels, respectively. H bonds are shown as grey dashed lines and their distances are shown in purple.

#### 2.5.2. Refinement of the neutron structure of *Prochlorococcus* iron binding FutA protein

Finally, we employed *REFMAC*5 for the refinement of a novel neutron structure. The marine cyanobacterium *Prochlorococcus* plays a significant role in global photosynthesis (Huston & Wolverton, 2009). However, its growth and productivity are constrained by the limited availability of iron. *Prochlorococcus* encodes the FutA protein that can accommodate binding of iron in either its ferric (Fe^3+^) or ferrous (Fe^2+^) state. The structure of FutA protein has been recently determined using a combination of structural biology techniques at room temperature, revealing the redox switch that allows the binding of both Fe oxidation states (Bolton *et al*., 2023).

The X-ray structure of FutA, determined at a resolution of 1.7 Å, shows that the iron binding site involves four tyrosine side chains (Tyr13, Tyr143, Tyr199, and Tyr200) and a solvent molecule, forming a trigonal bipyramidal coordination. The presence of Arg203 in the second coordination shell suggested the possibility of X-ray induced photoreduction of the iron centre, leading to a ferrous (Fe^2+^) binding state. To investigate the protonation of active site residues surrounding the iron, the neutron structure of FutA was determined at 2.1 Å resolution using ^1^H/D-exchanged crystals taking advantage of deuterium fraction refinement (Bolton *et al*., 2023). The final model is characterized by *R*_work_/*R*_free_ values of 18.2/25.0%. Data collection and refinement statistics are given in Table S2. The neutron structure reveals that the side chain of Arg103 is protonated and thus carries a positive charge, with all its exchangeable H atoms refining with deuterium fraction parameter values > 0.50 (Figure 5). Neutron maps suggest that the iron-coordinating residues Tyr13, Tyr143, Tyr199, and Tyr200 exist as tyrosinates. The H-omit map for the water molecule (W1) confirm its presence as neutral H_2_O, supported by the presence of positive peaks for two hydrogen atoms whose deuterium fraction values refine at 0.84 (H1) and 1.0 (H2) (Figure 5). In contrast to the room temperature X-ray structure, Arg203 is not involved in any interactions and does not contribute to the second coordination shell. Consequently, the Fe-binding site is composed of four negatively charged tyrosinates, a positively charged arginine in the second shell, and a neutral water (W1) suggesting that that this coordination cage neutralized ferric iron. This was further confirmed by the serial femtoseconds X-ray structure and electron paramagnetic resonance (EPR) measurements. Coordinates and structure factors of the neutron FutA have been deposited with the PDB under accession number 8OEN. This represents the first neutron structure refined and deposited within the PDB using *REFMAC5.* Further mechanistic information on FutA is discussed in a separate publication (Bolton *et al*., 2023).

**Figure 5.**
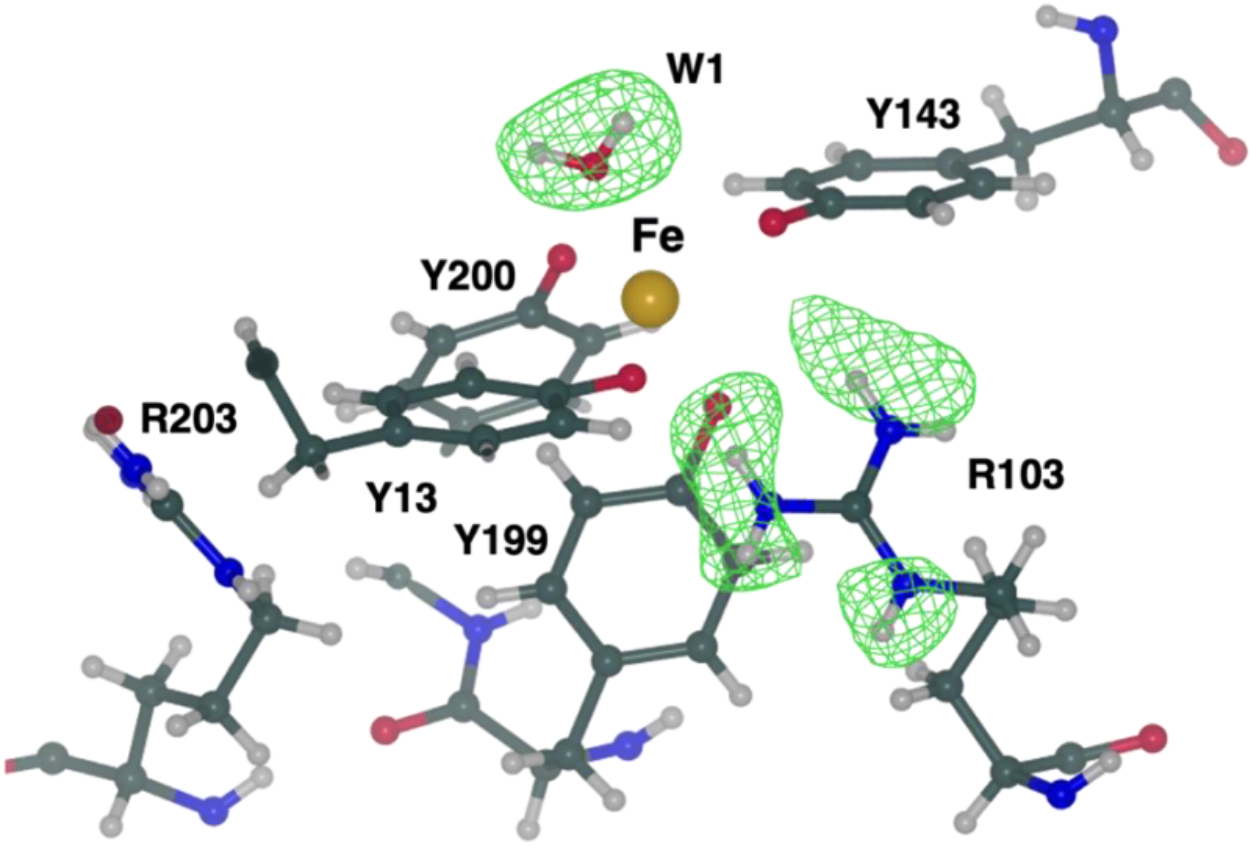
FutA (ferric state) determined by neutron diffraction at 2.1 Å resolution. The iron binding site is formed by four tyrosines (Tyr13, Tyr143, Tyr199, and Tyr200), a solvent molecule (W1) and Arg103 in the second coordination shell. The positive density (green mesh, *mF*_o_*−DF*_c_ omit map at +3.0σ level) indicates that these atoms have undergone ^1^H/D exchange and suggests that Arg103 is positively charged whilst W1 is neutral. The side chain of Arg203 is not oriented towards the binding site and does not engage in polar interactions. Nitrogen, carbon, and oxygen atoms are shown in blue, dark-green and red respectively. Iron is shown in gold and hydrogen atoms are in grey.

## 3. Conclusions and availability

Neutron crystallography offers a unique advantage in the determination of H atom positions, enabling the investigation of many biological processes. Despite its great potential in structural biology, the number of biological structures deposited to the PDB to date (25^th^ of July 2023) using neutron-only data (or joint neutron/X-ray) is extremely small (213), compared to those solved by X-ray crystallography (176,935), nuclear magnetic resonance (NMR) (14,034) and electron microscopy (EM) (16,239). This is due to technical limitations such as low neutron beam flux, long data collection times, and limited access to neutron beamlines. Nonetheless, recent advancements in instrumentation, experimental protocols, and computational tools have significantly advanced the field. As a result, the number of neutron structures deposited to the PDB has significantly increased in the last few years. The 2015-2022 period alone has seen the deposition of more than half the total neutron structures (130 out of 213), and this is likely to further accelerate in the coming years.

The *CCP*4 suite now provides tools for the refinement of macromolecular models using neutron diffraction data. Recent developments include the extension of the *CCP*4 Monomer Library by incorporating H atom nucleus distances. These restraints are required to ensure the correct H atom positions in neutron crystallographic refinement. Moreover, the inclusion of H nucleus positions holds potential for the further refinement of H atoms of cryo-EM models, as both H atom positions (electron and nucleus) contribute to the scattering. New features and refinement strategies have been implemented in *REFMAC*5 for the refinement of neutron models. Specifically, the introduction of the deuterium fraction parameter for H atoms. One of the benefits of this approach is that it generates models containing only H atoms, without any H/D or D sites. For each H atoms, the models incorporate a deuterium fraction parameter that indicates the level of deuteration in the sample. This results in clearer and more easily interpretable models, minimizing book-keeping errors that may arise when alternative conformations are present in the models. Re-refinement of neutron structures using *REFMAC5* has yielded *R*-factor values that are in line with the originally deposited values, including those obtained previously through joint neutron/X-ray techniques. Additionally, for certain neutron entries, the refinement process has led to improvements in model quality. Another valuable strategy is the use of external reference structure restraints during the refinement of models obtained by neutron diffraction. Incorporating restraints from X-ray reference structures has demonstrated an enhancement in the accuracy and reliability of neutron models, particularly in low resolution cases.

The ability to perform neutron crystallographic refinement using *REFMAC5* will be available in *CCP*4*i2* (Potterton *et al*., 2018) in an upcoming version of *CCP*4 that uses *Refmacat* instead of *REFMAC5*. This option can be enabled by selecting the appropriate diffraction experiment type (X-ray, Electron or Neutron) in the ’Advanced’ tab of the Refinement task interface, in which case the appropriate form factors and relevant default behaviours are used during model refinement. In Neutron mode, the graphical interface provides the ability to choose whether to refine all, only polar, or only rotatable polar H atom positions; to use H atom torsion angle restraints; to refine the H/D fraction for all H atoms (for perdeuterated crystals) or just polar H atoms (for H/D exchange experiments); and the choice of whether to reinitialise H/D fractions prior to refinement. Equivalent functionality will be available in an upcoming version of *CCP*4 *Cloud* (Krissinel *et al*., 2022). Relevant keywords and documentation for neutron crystallographic refinement will be available in the documentation section of the *CCP*4 website (https://www.ccp4.ac.uk/).

## Acknowledgements

The authors would like to thank Jake Grimmett, Toby Darling and Ivan Clayson for scientific computing resources. Paul Emsley is thanked for his assistance in preparing some of the figures. Rachel Bolton and Ivo Tews are thanked for useful conversations on the FutA structure.

## Funding information

LC is supported by an STFC/CCP4 PhD studentship (agreement No.7920 S2 2020 007) under the supervision of RAS and GNM. Part of this work was also supported by the BBSRC grant No. BB/P000169/1 awarded to RAS. KY, FL and GNM are supported by MRC grant No. MC_UP_A025_1012. RAN is supported by BBSRC grant No. BB/S007083/1.

## Footnotes

1 As a rule, when reference restraints or jelly-body refinement is used, more refinement cycles are needed to achieve the convergence.

## Notes

### Competing Interest Statement

The authors have declared no competing interest.

## References

1. Adams, P. D., Afonine, P. V., Bunkoczi, G., Chen, V. B., Davis, I. W., Echols, N., Headd, J. J., Hung, L. W., Kapral, G. J., Grosse-Kunstleve, R. W., McCoy, A. J., Moriarty, N. W., Oeffner, R., Read, R. J., Richardson, D. C., Richardson, J. S., Terwilliger, T. C. & Zwart, P. H. (2010). Acta Cryst. D 66, 213–221.

2. Adams, P. D., Mustyakimov, M., Afonine, P. V. & Langan, P. (2009). Acta Crystallographica Section D: Biological Crystallography 65, 567–573.

3. Afonine, P. V., Grosse-Kunstleve, R. W., Echols, N., Headd, J. J., Moriarty, N. W., Mustyakimov, M., Terwilliger, T. C., Urzhumtsev, A., Zwart, P. H. & Adams, P. D. (2012). Acta Cryst. D 68, 352–367.

4. Afonine, P. V., Mustyakimov, M., Grosse-Kunstleve, R. W., Moriarty, N. W., Langan, P. & Adams, P. D. (2010). Acta Cryst. D 66, 1153–1163.

5. Afonine, P. V., Poon, B. K., Read, R. J., Sobolev, O. V., Terwilliger, T. C., Urzhumtsev, A. & Adams, P. D. (2018). Acta Crystallogr D Struct Biol 74, 531–544.

6. Agirre, J., Atanasova, M., Bagdonas, H., Ballard, C. B., Basle, A., Beilsten-Edmands, J., Borges, R. J., Brown, D. G., Burgos-Marmol, J. J., Berrisford, J. M., Bond, P. S., Caballero, I., Catapano, L., Chojnowski, G., Cook, A. G., Cowtan, K. D., Croll, T. I., Debreczeni, J. E., Devenish, N. E., Dodson, E. J., Drevon, T. R., Emsley, P., Evans, G., Evans, P. R., Fando, M., Foadi, J., Fuentes-Montero, L., Garman, E. F., Gerstel, M., Gildea, R. J., Hatti, K., Hekkelman, M. L., Heuser, P., Hoh, S. W., Hough, M. A., Jenkins, H. T., Jimenez, E., Joosten, R. P., Keegan, R. M., Keep, N., Krissinel, E. B., Kolenko, P., Kovalevskiy, O., Lamzin, V. S., Lawson, D. M., Lebedev, A. A., Leslie, A. G. W., Lohkamp, B., Long, F., Maly, M., McCoy, A. J., McNicholas, S. J., Medina, A., Millan, C., Murray, J. W., Murshudov, G. N., Nicholls, R. A., Noble, M. E. M., Oeffner, R., Pannu, N. S., Parkhurst, J. M., Pearce, N., Pereira, J., Perrakis, A., Powell, H. R., Read, R. J., Rigden, D. J., Rochira, W., Sammito, M., Sanchez Rodriguez, F., Sheldrick, G. M., Shelley, K. L., Simkovic, F., Simpkin, A. J., Skubak, P., Sobolev, E., Steiner, R. A., Stevenson, K., Tews, I., Thomas, J. M. H., Thorn, A., Valls, J. T., Uski, V., Uson, I., Vagin, A., Velankar, S., Vollmar, M., Walden, H., Waterman, D., Wilson, K. S., Winn, M. D., Winter, G., Wojdyr, M. & Yamashita, K. (2023). Acta Crystallographica Section D 79, 449–461.

7. Ahmed, H. U., Blakeley, M. P., Cianci, M., Cruickshank, D. W. J., Hubbard, J. A. & Helliwell, J. R. (2007). Acta Crystallographica Section D 63, 906–922.

8. Allen, F., Kennard, O., Watson, D., Brammer, L., Orpen, A. & Taylor, R. (1992). pp. 685-706. Dordrecht: Kluwer Academic Publishers.

9. Allen, F. H. & Bruno, I. J. (2010). Acta Cryst. B 66, 380–386.

10. Allen, F. H., Kennard, O., Watson, D. G., Brammer, L., Orpen, A. G. & Taylor, R. (1987). Journal of the Chemical Society, Perkin Transactions 2 S1–S19.

11. Baker, L. A. & Rubinstein, J. L. (2010). Vol. 481, Methods Enzymol., edited by G. J. Jensen, pp. 371-388. Academic Press.

12. Berman, H. M., Westbrook, J., Feng, Z., Gilliland, G., Bhat, T. N., Weissig, H., Shindyalov, I. N. & Bourne, P. E. (2000). Nucleic Acids Res. 28, 235–242.

13. Blakeley, M. P. & Podjarny, A. D. (2018). Emerg Top Life Sci 2, 39–55.

14. Bolton, R., Machelett, M. M., Stubbs, J., Axford, D., Caramello, N., Catapano, L., Malý, M., Rodrigues, M. J., Cordery, C., Tizzard, G. J., MacMillan, F., Engilberge, S., Stetten, D. v., Tosha, T., Sugimoto, H., Worrall, J. A. R., Webb, J. S., Zubkov, M., Coles, S., Mathieu, E., Steiner, R. A., Murshudov, G., Schrader, T. E., Orville, A. M., Royant, A., Evans, G., Hough, M. A., Owen, R. L. & Tews, I. (2023). bioRxiv 2023.2005.2023.541926.

15. Brünger, A. T., Adams, P. D., Clore, G. M., DeLano, W. L., Gros, P., Grosse-Kunstleve, R. W., Jiang, J.-S., Kuszewski, J., Nilges, M. & Pannu, N. S. (1998). Acta Crystallographica Section D: Biological Crystallography 54, 905–921.

16. Bruno, I. J., Cole, J. C., Edgington, P. R., Kessler, M., Macrae, C. F., McCabe, P., Pearson, J. & Taylor, R. (2002). Acta Cryst. B 58, 389–397.

17. Budayova-Spano, M., Koruza, K. & Fisher, Z. (2020). Methods Enzymol. 634, 21–46.

18. Bui, S., von Stetten, D., Jambrina, P. G., Prange, T., Colloc’h, N., de Sanctis, D., Royant, A., Rosta, E. & Steiner, R. A. (2014). Angew. Chem. Int. Ed. Engl. 53, 13710–13714.

19. Chatake, T., Mizuno, N., Voordouw, G., Higuchi, Y., Arai, S., Tanaka, I. & Niimura, N. (2003). Acta Crystallographica Section D 59, 2306–2309.

20. Chen, J. C.-H., Fisher, Z., Kovalevsky, A. Y., Mustyakimov, M., Hanson, B. L., Zhurov, V. V. & Langan, P. (2012). Acta Crystallographica Section F 68, 119–123.

21. Clabbers, M. T. B. & Abrahams, J. P. (2018). Crystallography Reviews 24, 176–204.

22. Clabbers, M. T. B., Martynowycz, M. W., Hattne, J. & Gonen, T. (2022). J Struct Biol X 6, 100078.

23. Combs, J. E., Andring, J. T. & McKenna, R. (2020). Methods Enzymol. 634, 281–309.

24. Coppens, P. (1997). X-ray charge densities and chemical bonding. International Union of Crystallography.

25. Coppens, P., Boehme, R., Price, P. & Stevens, E. (1981). Acta Crystallographica Section A: Crystal Physics, Diffraction, Theoretical and General Crystallography 37, 857–863.

26. Diamond, R. (1971). Acta Crystallographica Section A: Crystal Physics, Diffraction, Theoretical and General Crystallography 27, 436–452.

27. Engler, N., Ostermann, A., Niimura, N. & Parak, F. G. (2003). Proc. Natl. Acad. Sci. U. S. A. 100, 10243–10248.

28. Fermi, E. & Marshall, L. (1947). Physical Review 72, 1139–1146.

29. Fisher, S., Blakeley, M., Cianci, M., McSweeney, S. & Helliwell, J. (2012). Acta Crystallographica Section D: Biological Crystallography 68, 800–809.

30. Fisher, S. J., Blakeley, M. P., Howard, E. I., Petit-Haertlein, I., Haertlein, M., Mitschler, A., Cousido-Siah, A., Salvay, A. G., Popov, A., Muller-Dieckmann, C., Petrova, T. & Podjarny, A. (2014). Acta Cryst. D 70, 3266–3272.

31. Fisher, S. J., Wilkinson, J., Henchman, R. H. & Helliwell, J. R. (2009). Crystallography Reviews 15, 231–259.

32. French, S. & Wilson, K. (1978). Acta Crystallographica Section A 34, 517–525.

33. Gardberg, A. S., Del Castillo, A. R., Weiss, K. L., Meilleur, F., Blakeley, M. P. & Myles, D. A. (2010). Acta Cryst. D 66, 558–567.

34. Garman, E. F. (2010). Acta Cryst. D 66, 339–351.

35. Grazulis, S., Daskevic, A., Merkys, A., Chateigner, D., Lutterotti, L., Quiros, M., Serebryanaya, N. R., Moeck, P., Downs, R. T. & Le Bail, A. (2012). Nucleic Acids Res. 40, D420–427.

36. Groom, C. R., Bruno, I. J., Lightfoot, M. P. & Ward, S. C. (2016). Acta Crystallogr B Struct Sci Cryst Eng Mater 72, 171–179.

37. Gruene, T., Hahn, H. W., Luebben, A. V., Meilleur, F. & Sheldrick, G. M. (2014). J Appl Crystallogr 47, 462–466.

38. Guzenko, D., Burley, S. K. & Duarte, J. M. (2020). PLoS Comput. Biol. 16, e1007970.

39. Headd, J. J., Echols, N., Afonine, P. V., Grosse-Kunstleve, R. W., Chen, V. B., Moriarty, N. W., Richardson, D. C., Richardson, J. S. & Adams, P.D. (2012). Acta Crystallographica Section D: Biological Crystallography 68, 381–390.

40. Howard, E. I., Sanishvili, R., Cachau, R. E., Mitschler, A., Chevrier, B., Barth, P., Lamour, V., Van Zandt, M., Sibley, E., Bon, C., Moras, D., Schneider, T. R., Joachimiak, A. & Podjarny, A. (2004). Proteins: Structure, Function, and Bioinformatics 55, 792–804.

41. Huston, M. A. & Wolverton, S. (2009). Ecol. Monogr. 79, 343–377.

42. Jack, A. & Levitt, M. (1978). Acta Crystallographica Section A: Crystal Physics, Diffraction, Theoretical and General Crystallography 34, 931–935.

43. Kahn, K. (1999). Bioorg. Chem. 27, 351–362.

44. Kahn, K., Serfozo, P. & Tipton, P. A. (1997). J. Am. Chem. Soc. 119, 5435–5442.

45. Kneller, D. W., Li, H., Phillips, G., Weiss, K. L., Zhang, Q., Arnould, M. A., Jonsson, C. B., Surendranathan, S., Parvathareddy, J., Blakeley, M. P., Coates, L., Louis, J. M., Bonnesen, P. V. & Kovalevsky, A. (2022). Nature Communications 13, 2268.

46. Konnert, J. H. & Hendrickson, W. A. (1980). Acta Crystallographica Section A: Crystal Physics, Diffraction, Theoretical and General Crystallography 36, 344–350.

47. Kossiakoff, A. A. (1984). Neutrons in Biology, edited by B. P. Schoenborn, pp. 281-304. Boston, MA: Springer US.

48. Kovalevsky, A., Gerlits, O., Beltran, K., Weiss, K. L., Keen, D. A., Blakeley, M. P., Louis, J. M. & Weber, I. T. (2020). pp. 257–279. Elsevier.

49. Krissinel, E., Lebedev, A. A., Uski, V., Ballard, C. B., Keegan, R. M., Kovalevskiy, O., Nicholls, R. A., Pannu, N. S., Skubak, P., Berrisford, J., Fando, M., Lohkamp, B., Wojdyr, M., Simpkin, A. J., Thomas, J. M. H., Oliver, C., Vonrhein, C., Chojnowski, G., Basle, A., Purkiss, A., Isupov, M. N., McNicholas, S., Lowe, E., Trivino, J.,

50. Cowtan, K., Agirre, J., Rigden, D. J., Uson, I., Lamzin, V., Tews, I., Bricogne, G., Leslie, A. G. W. & Brown, D. G. (2022). Acta Crystallographica Section D 78, 1079–1089.

51. Lawson, C. L., Patwardhan, A., Baker, M. L., Hryc, C., Garcia, E. S., Hudson, B. P., Lagerstedt, I., Ludtke, S. J., Pintilie, G., Sala, R., Westbrook, J. D., Berman, H. M., Kleywegt, G. J. & Chiu, W. (2015). Nucleic Acids Res. 44, D396–D403.

52. Liebschner, D., Afonine, P. V., Moriarty, N. W., Langan, P. & Adams, P. D. (2018). Acta Crystallographica Section D 74, 800–813.

53. Logan, D. T. (2020). Vol. 634, Methods Enzymol., edited by P. C. E. Moody, pp. 201-224. Academic Press.

54. Long, F., Nicholls, R. A., Emsley, P., Graaeulis, S., Merkys, A., Vaitkus, A. & Murshudov, G. N. (2017). Acta Crystallogr D Struct Biol 73, 112–122.

55. Macrae, C. F., Sovago, I., Cottrell, S. J., Galek, P. T. A., McCabe, P., Pidcock, E., Platings, M., Shields, G. P., Stevens, J. S., Towler, M. & Wood, P. A. (2020). J Appl Crystallogr 53, 226–235.

56. Maki-Yonekura, S., Kawakami, K., Takaba, K., Hamaguchi, T. & Yonekura, K. (2023). Communications Chemistry 6, 98.

57. McGregor, L., Foldes, T., Bui, S., Moulin, M., Coquelle, N., Blakeley, M. P., Rosta, E. & Steiner, R. A. (2021). IUCrJ 8, 46–59.

58. Meilleur, F., Weiss, K. L. & Myles, D. A. A. (2009). Micro and Nano Technologies in Bioanalysis: Methods and Protocols, edited by R. S. Foote & J. W. Lee, pp. 281-292. Totowa, NJ: Humana Press.

59. Murshudov, G. N., Skubak, P., Lebedev, A. A., Pannu, N. S., Steiner, R. A., Nicholls, R. A., Winn, M. D., Long, F. & Vagin, A. A. (2011). Acta Cryst. D 67, 355–367.

60. Nakane, T., Kotecha, A., Sente, A., McMullan, G., Masiulis, S., Brown, P. M. G. E., Grigoras, I. T., Malinauskaite, L., Malinauskas, T., Miehling, J., Yu, L., Karia, D., Pechnikova, E. V., De Jong, E., Keizer, J., Bischoff, M., McCormack, J., Tiemeijer, P., Hardwick, S. W., Chirgadze, D. Y., Murshudov, G., Aricescu, A. R. & Scheres, S. H. W. (2020).

61. Nicholls, R. A., Fischer, M., McNicholas, S. & Murshudov, G. N. (2014). Acta Cryst. D 70, 2487–2499.

62. Nicholls, R. A., Joosten, R. P., Long, F., Wojdyr, M., Lebedev, A., Krissinel, E., Catapano, L., Fischer, M., Emsley, P. & Murshudov, G. N. (2021a). Acta Crystallogr D Struct Biol 77, 712–726.

63. Nicholls, R. A., Long, F. & Murshudov, G. N. (2012). Acta Cryst. D 68, 404–417.

64. Nicholls, R. A., Tykac, M., Kovalevskiy, O. & Murshudov, G. N. (2018). Acta Crystallographica Section D: Structural Biology 74, 492–505.

65. Nicholls, R. A., Wojdyr, M., Joosten, R. P., Catapano, L., Long, F., Fischer, M., Emsley, P. & Murshudov, G. N. (2021b). Acta Crystallogr D Struct Biol 77, 727–745.

66. Niimura, N., Chatake, T., Kurihara, K. & Maeda, M. (2004). Cell Biochem. Biophys. 40, 351–370.

67. Niimura, N. & Podjarny, A. (2011). Neutron protein crystallography: hydrogen, protons, and hydration in bio-macromolecules. Oxford University Press.

68. Oksanen, E., Chen, J. C. H. & Fisher, S. Z. (2017). Molecules 22, 596.

69. Orpen, A. G., Brammer, L., Allen, F. H., Kennard, O., Watson, D. G. & Taylor, R. (1989). Journal of the Chemical Society, Dalton Transactions S1–S83.

70. Petrova, T. & Podjarny, A. (2004). Reports on Progress in Physics 67, 1565–1605.

71. Pierce, J., Cuneo, M. J., Jennings, A., Li, L., Meilleur, F., Zhao, J. & Myles, D. A. A. (2020). Methods Enzymol. 634, 153–175.

72. Potterton, L., Agirre, J., Ballard, C., Cowtan, K., Dodson, E., Evans, P. R., Jenkins, H. T., Keegan, R., Krissinel, E., Stevenson, K., Lebedev, A., McNicholas, S. J., Nicholls, R. A., Noble, M., Pannu, N. S., Roth, C., Sheldrick, G., Skubak, P., Turkenburg, J., Uski, V., von Delft, F., Waterman, D., Wilson, K., Winn, M. & Wojdyr, M. (2018). Acta Crystallogr D Struct Biol 74, 68–84.

73. Schmidt, M. W., Baldridge, K. K., Boatz, J. A., Elbert, S. T., Gordon, M. S., Jensen, J. H., Koseki, S., Matsunaga, N., Nguyen, K. A., Su, S. J., Windus, T. L., Dupuis, M. & Montgomery, J. A. (1993). J. Comput. Chem. 14, 1347–1363.

74. Schröder, G. F., Levitt, M. & Brunger, A. T. (2014). Acta Cryst. D 70, 2241–2255.

75. Sears, V. F. (1992). Neutron News 3, 26–37.

76. Shabalin, I. G., Porebski, P. J. & Minor, W. (2018). Crystallogr Rev 24, 236–262.

77. Sheldrick, G. M. (2015). Acta Crystallogr C Struct Chem 71, 3–8.

78. Sheldrick, G. M. & Schneider, T. R. (1997). Methods Enzymol. 277, 319–343.

79. Shu, F., Ramakrishnan, V. & Schoenborn, B. P. (2000). Proceedings of the National Academy of Sciences 97, 3872–3877.

80. Smart, O. S., Womack, T. O., Flensburg, C., Keller, P., Paciorek, W., Sharff, A., Vonrhein, C. & Bricogne, G. (2012). Acta Crystallographica Section D 68, 368–380.

81. Tronrud, D. E. (2004). Acta Cryst. D 60, 2156–2168.

82. Vagin, A. A., Steiner, R. A., Lebedev, A. A., Potterton, L., McNicholas, S., Long, F. & Murshudov, G. N. (2004). Acta Cryst. D 60, 2184–2195.

83. van Beusekom, B., Joosten, K., Hekkelman, M. L., Joosten, R. P. & Perrakis, A. (2018). IUCrJ 5, 585–594.

84. Wan, Q., Parks, J. M., Hanson, B. L., Fisher, S. Z., Ostermann, A., Schrader, T. E., Graham, D. E., Coates, L., Langan, P. & Kovalevsky, A. (2015). Proceedings of the National Academy of Sciences 112, 12384–12389.

85. Waser, J. (1963). Acta Cryst. 16, 1091–1094.

86. Wei, D., Huang, X., Qiao, Y., Rao, J., Wang, L., Liao, F. & Zhan, C.-G. (2017). ACS catalysis 7, 4623–4636.

87. Wishart, D. S., Feunang, Y. D., Guo, A. C., Lo, E. J., Marcu, A., Grant, J. R., Sajed, T., Johnson, D., Li, C., Sayeeda, Z., Assempour, N., Iynkkaran, I., Liu, Y., Maciejewski, A., Gale, N., Wilson, A., Chin, L., Cummings, R., Le, D., Pon, A., Knox, C. & Wilson, M. (2018). Nucleic Acids Res. 46, D1074–D1082.

88. Wojdyr, M. (2022). Journal of Open Source Software 7, 4200.

89. Yamashita, K., Palmer, C. M., Burnley, T. & Murshudov, G. N. (2021). Acta Cryst. D 77, 1282–1291.

90. Yamashita, K., Wojdyr, M., Long, F., Nicholls, R. A. & Murshudov, G. N. (2023). Acta Crystallogr D Struct Biol 79, 368–373.

91. Yip, K. M., Fischer, N., Paknia, E., Chari, A. & Stark, H. (2020). Nature 587, 157–161.

