## Supplementary material for "Neutron crystallographic refinement with *REFMAC*5 of the *CCP*4 suite": Table S1 and Table S2

Supporting information

Table S1 Neutron data collection and refinement statistics for UOX:8AZA complex

| Data collection statistics | | |
| --- | --- | --- |
| Space group | | *I*222 |
| Temperature (°C) | | 20 |
| Wavelength (Å) | | 3.12 – 4.20 |
| Unit Cell (a, b, c; Å) | | 80.6, 96.1, 105.4 |
| Resolution range (Å)^1^ | | 40.00 – 2.10 (2.15 – 2.10) |
| CC ½ (%)^1^ | | 98.7 (82.6) |
| < I/σ (I)^1^> | | 5.8 (2.9) |
| Completeness (%)^1^ | | 78.62 (65.89) |
| Unique reflections^1^ | | 18,131(1099) |
| Refinement statistics | | |
| Resolution (Å) | | 40.00 – 2.10 |
| *R*_work_/*R*_free_ (%) | | 17.0 / 26.0 |
| No. of reflections all/free | | 18,131/500 |
| No. of | |  |
| Protein residues | | 295 |
| Solvent molecules  Ligands  Ions | | 350  1  1 |
| Avarage *B* factors (Å^2^) | |  |
|  | Protein | 45.18 |
|  | Solvent | 55.21 |
|  | Ligand | 36.23 |
|  | Ion | 44.00 |
| r.m.s. deviations | |  |
|  | Bond lengths (Å) | 0.006 |
|  | Bond angles (°) | 0.961 |

^1^High resolution statistics in parentheses

Table S2 Neutron data collection and refinement statistics for FutA structure

| Data collection statistics | | |
| --- | --- | --- |
| Space group | | *P*2_1_ |
| Temperature (°C) | | 21 |
| Wavelength (Å) | | 3.10 |
| Unit Cell (a, b, c; Å) | | 39.5, 78.3, 47.9 |
| β angle (°) | | 97.4 |
| Resolution range (Å)^1^ | | 24.94 – 2.1 (2.18 – 2.1) |
| CC ½ (%)^1^ | | 97.1 (68.1) |
| < I/σ (I)^1^> | | 4.8 (2.0) |
| Completeness (%)^1^ | | 81.0 (52.6) |
| Multiplicity^1^ | | 1.9 (1.1) |
| Unique reflections^1^ | | 13,731 (890) |
| Refinement statistics | | |
| PDB code | | 8OEN |
| Resolution (Å) | | 24.95 – 2.1 |
| *R*_work_/*R*_free_ (%) | | 18/2 / 25.0 |
| No. of reflections all/free | | 13,731 / 698 |
| No. of | |  |
|  | Protein residues | 314 |
|  | Solvent molecules | 96 |
|  | Ions | 1 |
| Avarage *B* factors (Å^2^) | |  |
|  | Protein | 22.31 |
|  | Solvent | 13.41 |
| r.m.s. deviations | |  |
|  | Bond lengths (Å) | 0.005 |
|  | Bond angles (°) | 1.142 |

^1^High resolution statistics in parentheses
